# Interplay of *Mycobacterium abscessus* and *Pseudomonas aeruginosa* in coinfection: Biofilm Dynamics and Host Immune Response

**DOI:** 10.1101/2024.01.22.576702

**Authors:** Víctor Campo-Pérez, Esther Julián, Eduard Torrents

**Author notes:** Corresponding authors: Dr. Esther Julián, Dr. Eduard Torrents /.

## Abstract

The incidence of infection by nontuberculous mycobacteria, mainly *Mycobacterium abscessus*, in patients with cystic fibrosis and other chronic pulmonary illnesses is increasing, translating into an acceleration in the decline of lung function. In most cases, *M. abscessus* coinfects with *Pseudomonas aeruginosa*, the most common pathogen in these chronic diseases. However, it is unknown how these two bacterial species interact when coinfecting. This study aims to explore the behavior of both species in three relevant pathogenic settings: dual-species biofilm development using a recently developed method to monitor individual species in dual-species biofilms; coinfection in bronchial epithelial cells using *in vitro* assays; and *in vivo* coinfection using the *Galleria mellonella* model. The results demonstrate the capability of both species to form stable mixed biofilms and to reciprocally inhibit single-biofilm progression. Coinfections in bronchial epithelial cells were correlated with significantly decreased cell viability, while in *G. mellonella,* coinfections induced lower survival rates than individual infections. Outstandingly, the analysis of the immune response triggered by each bacterium in bronchial epithelial cell assays and *G. mellonella* larvae revealed that *P. aeruginosa* induces the overexpression of proinflammatory and melanization cascade responses, respectively. In contrast, *M. abscessus* and *P. aeruginosa* coinfection significantly inhibited the immune response in both models, resulting in worse consequences for the host than those generated by single *P. aeruginosa* infection. Overall, the presence of *M. abscessus* produces a decline in the immune responses that worsens the infection and compromises the host.

**Importance:** The appearance of bacterial infections in the respiratory tract of patients with chronic respiratory diseases suppose a serious and difficult to treat health problem. This complication is exacerbated by the increase resistance against antibiotics generated by pathogenic microorganisms. The most common and virulent pathogenic bacteria reported in the respiratory airway is *Pseudomonas aeruginosa*. It is a Gram-negative, ubiquitous, and intrinsic resistant to antibiotics bacteria. However, the incidence of a rapidly growing, multi-drug resistant mycobacteria; *Mycobacterium abscessus*, is growing worldwide. The pulmonary coinfection by both pathogens is directly related with higher rates of morbidity and mortality of patients. The significance of our research is characterizing the behavior of these two pathogens when they coinfects together, exploring the immune response triggered by the host and its impact in the survival. The purpose is enhancing the limited understanding we have of this clinically relevant coinfection to favor the development of new effective treatments.

## Introduction

Cystic fibrosis (CF) is an autosomal recessive inherited condition caused by the presence of mutations in the gene encoding the cystic fibrosis transmembrane conductance regulator (CFTR) protein (1). This protein acts as a transporter channel that conducts chloride ions (Cl^-^) across epithelial cell membranes to maintain the balance of salt and water on many surfaces in the body and is especially relevant in the respiratory system airways (2). A malfunction in CFTR protein prevents proper hydration of the cellular surface, which leads to the loss of the mucus covering the cells, becoming thick and sticky (3). These characteristics of the mucus in CF patients hinder the primary innate defense mechanism of the lung, mucociliary clearance (4), leading to an increased risk of infections from different pathogens, especially *Pseudomonas aeruginosa*, *Staphylococcus aureus*, *Burkholderia spp.* and nontuberculous *Mycobacterium spp.* (NTM), which are associated with a high risk of pulmonary compromise and lead to chronic infections (5). In fact, lung tract infections in CF patients are complex and polymicrobial, capable of developing intermicrobial and host‒pathogen interactions impacting clinically relevant features such as virulence, persistent colonization of the airways, and antimicrobial recalcitrance (6, 7). Although CF polymicrobial infections are highly individualized in each patient, *S. aureus* and *P. aeruginosa* dominate in children and adult microbial communities, respectively. In this sense, the coinfection by these two dominant pathogens or one of them with other species negatively affect respiratory function compared to single infections (7).

*P. aeruginosa* is a ubiquitous gram-negative opportunistic pathogen that is metabolically flexible and considered the most relevant bacterium associated with CF due to its high prevalence and pathogenesis. Due to its ability to adapt to the CF pulmonary environment, *P. aeruginosa* is also the primary pathogen present in advanced CF lung disease (8). *P. aeruginosa* PAO1 is the reference laboratory strain for research and PAET1 strain is a clinical isolate from a cystic fibrosis chronically infected patient. *P. aeruginosa* develops biofilms on the thick and dehydrated mucoid surface, which represents a difficult barrier to antibiotic penetration and to host innate immune effectors, in CF (9). These biofilms cause chronic infections through higher antibiotic tolerance, resistance to phagocytosis, and immune-mediated chronic inflammation (10). Furthermore, biofilms are usually polymicrobial and are related to an exacerbated inflammatory immune response that seriously compromises respiratory function, producing the most extensive lung lesions (11).

A significant increase in the incidence of infections caused by NTM in CF patients has been recently reported (12, 13), with fast-growing *Mycobacterium abscessus* being the most prominent and worrisome pathogen of this genus (14). *M. abscessus* infections are associated with an increase in morbidity and mortality due to the long duration of treatment related to significant adverse effects, its multiresistance to antimicrobials, and consequently, the low success rate in its elimination from the respiratory tract (15). Another relevant element in *M. abscessus* pathogenicity is its ability to modify its cell wall, which allows the identification of two different morphotypes in this species: smooth (S), which contains abundant glycopeptidolipids (GPLs) in the outer layer, and rough (R), which is devoid of GPLs (16). Similar to *P. aeruginosa*, *M. abscessus* also has a biofilm-forming capacity, making treatment even more difficult. Remarkably, 58-78% of patients with *M. abscessus* infection are also infected by *P. aeruginosa* (17). The colonization of the lung epithelium begins with S morphotypes, which progressively switch to R mutants, giving a more aggressive, invasive pulmonary infection (18). In this sense, biofilms developed by each morphotype of *M. abscessus* present different mechanical properties, with R biofilms having higher rigidity than S biofilms. Interestingly, *M. abscessus* R/S biofilms present more resistance to mechanical forms of clearance from the lung than *P. aeruginosa* biofilms (19). Furthermore, *M. abscessus* biofilm bacilli change the mycolic acid proportions and produce a lipid-relevant extracellular matrix with other components, such as carbohydrates, proteins, or extracellular DNA (20).

Coinfections of *P. aeruginosa* and *M. abscessus* in CF have become clinically more relevant over the last two decades; the incidence of NTM infections among CF patients has risen from 3.3% to 22.6%, resulting in an increase in morbidity and mortality (16). In 16% to 68% of NTM-positive sputum cultures, the *M. abscessus* complex is present, and its prevalence is increasing (12). Furthermore, the presence of *M. abscessus* lung disease has been considered a strong relative contraindication to lung transplantation (21), which can extend and improve the quality of life in properly CF patients, increasing their mean life expectancy by 7.4 years (22).

Despite the challenges stated above, studies focused on the interaction between both species are scarce. It is therefore crucial to further elucidate and characterize the *P. aeruginosa*-*M. abscessus* coinfections. For this reason, the objective of the present study is to unravel the interaction mechanisms between both bacteria, focusing on relevant aspects of pathogenicity: dual-species biofilm development using recently developed coculture techniques, including S and R *M. abscessus* morphotypes, and the viability outcome and the analysis of the triggered immune response by each species in individual and coinfection experiments using *in vitro* and *in vivo* models. A better understanding of the behavior between *P. aeruginosa* and *M. abscessus* would provide us with tools to develop efficient treatment strategies against these pathogens in CF patients.

## Results

### M. abscessus reduces the progression of P. aeruginosa biofilms in vitro

*In vitro* biofilm development assays (see Material and Methods) showed that the addition of *M. abscessus* R or S to mature *P. aeruginosa* biofilms limits their growth capacity (**Figure 1A**). This inhibitory effect was observed for both strains of *P. aeruginosa* PAO1 and PAET1 biofilms. Specifically, the presence of *M. abscessus* (R and S) resulted in a growth reduction of approximately 20% for PAO1 and 30% for PAET1 biofilms (**Figure 1A**) compared with the nontreated *P. aeruginosa* biofilms (represented as 100% growth). Nevertheless, the highest rate of growth inhibition was reported with *M. abscessus* S at 10^4^ cfus/well, with approximately 40% PAO1 biofilm reduction. The addition of heat-killed *M. abscessus* or bacterial culture supernatants had no inhibitory effect on the growth of *P. aeruginosa* mature biofilms (**Figure 1A**). Similarly, the use of *E. coli* (gram-negative) or *B. thuringiensis* (gram-positive) did not have a growth-limiting effect on *P. aeruginosa* biofilms and even, in the case of heat-killed *E. coli* and supernatant from *E. coli* cultures, induced *P. aeruginosa* PAO1 biofilm growth (**Figure 1A**).

**Figure 1:**
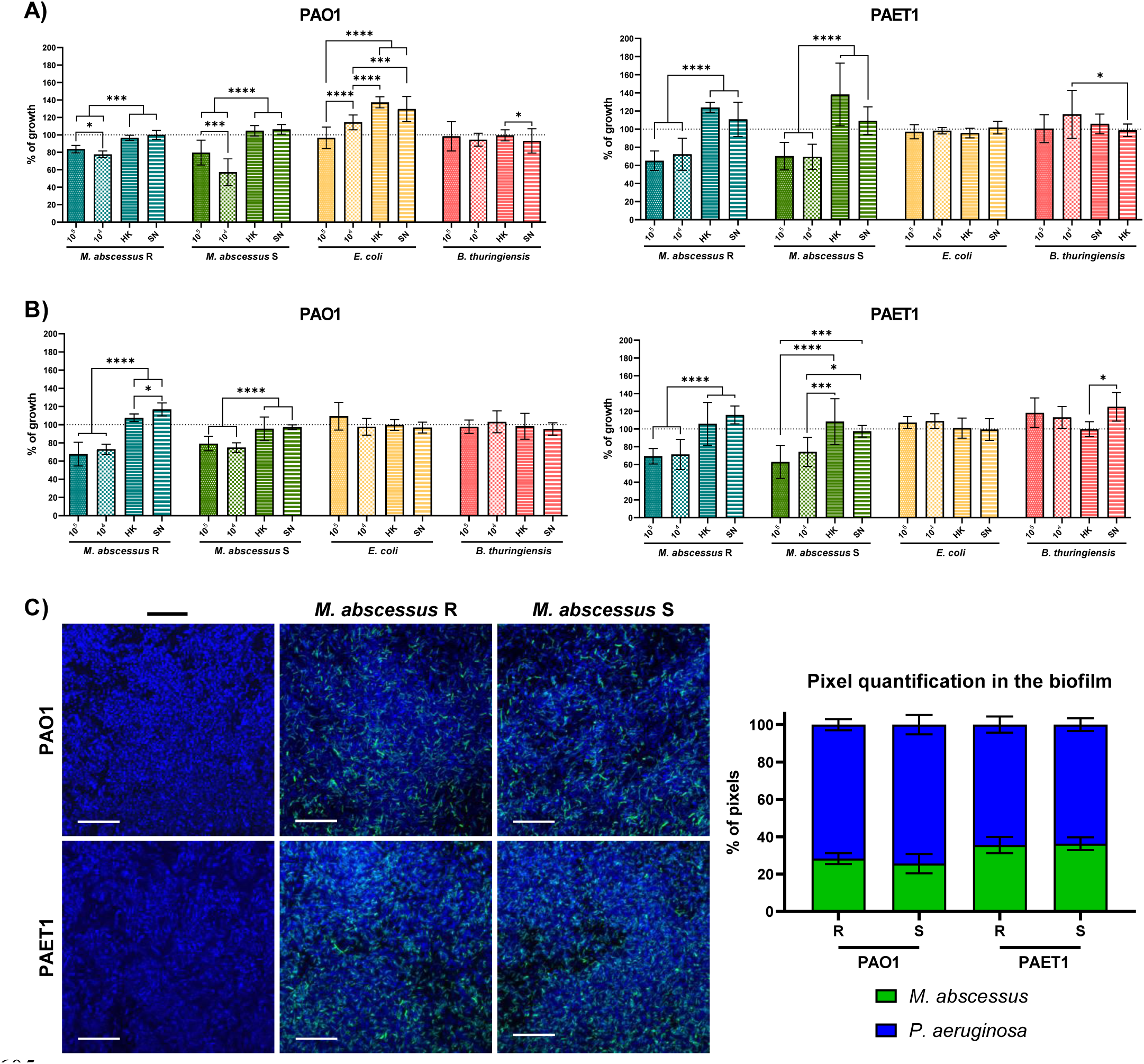
Coculture of *P. aeruginosa* PAO1 and PAET1 biofilms with different bacteria. **A**) Growth of well-developed mature *P. aeruginosa* biofilm when different bacterial cultures (live and heat-killed) or bacterial culture supernatants were added. **B)** Growth of *P. aeruginosa* in coculture biofilms. The values correspond to the luciferase expression (*lux*) of the *P. aeruginosa* strains generated in each condition, normalized based on the average expression reported in control wells (marked with a dashed line as 100% *P. aeruginosa* growth). The effects of *M. abscessus* R and S morphotypes, *E. coli* MG1655 and *B. thuringiensis* on both *P. aeruginosa* strains were tested as well as the supernatant (SN) and heat-killed (HK) form of these bacterial cultures. Data are presented as the mean values ± SDs of at least three independent experiments including six replicates in each experimental condition; significant differences were determined using one-way ANOVA and Tukey’s multiple comparisons tests; ****, *p* < 0.0001; ***, *p* < 0.001; **, *p* < 0.01; *, *p* < 0.05. **C)** Representative confocal microscopy images of coculture biofilms and pixel color quantification of the images. *P. aeruginosa* PAO1 and PAET1 biofilms were grown for 72 h, and *M. abscessus* R and S were inoculated for an additional 24 h. Z-stack compositions of 96 h-old *P. aeruginosa* PAO1 and PAET1 biofilms in combination with *M. abscessus* R and S morphotypes. DAPI stained all cells blue, and green fluorescent protein (GFP) of plasmid pETS218 allowed mycobacterial cells to be detected in green. Scale bars correspond to 20 µm. Pixel quantifications are represented as the mean values ± SDs from three representative images of each condition.

When *P. aeruginosa* strains were cocultured simultaneously with all bacteria tested, the results were comparable with the previous assay results but with slight inhibition percentage changes in some cases (**Figure 1B**). *P. aeruginosa* PAO1 cocultured with the *M. abscessus* R morphotype showed a 30% reduction in PAO1 biofilm compared to the control, while with the S morphotype, the reduction in PAO1 biofilm was only approximately 20%. In *P. aeruginosa* PAET1 cocultures, the results using *M. abscessus* R or S were quite similar, with an approximately 30% decrease in PAET1 biofilm compared to the control (growth of *P. aeruginosa* alone) (**Figure 1B**). As previously shown in Figure 1A, the addition of *E. coli* and *B. thuringiensis* and the use of heat-killed cultures or supernatants did not have an inhibitory effect on the *P aeruginosa* PAO1 and PAET1 biofilms (**Figure 1B**). Also, it was observed that PAET1 biofilms progression was highly inhibited in coculture with *M. abscessus* (R and S) than those of PAO1 in all the biofilm assays tested.

After observation of a clear *P. aeruginosa* biofilm reduction with the addition of *M. abscessus*, either added in a mature biofilm or when simultaneously grown in a dual-species biofilm, the structure and composition of coculture biofilms were analyzed by confocal microscopy. The images showed that after 24 h of coculture, *M. abscessus* was established in a *P. aeruginosa* mature biofilm, demonstrating the capacity to settle and simultaneously grow under these conditions (**Figure 1C**). The presence of *P. aeruginosa* cells (in blue) was clearly higher, but *M. abscessus* bacilli (in green) were clearly observed in both *P. aeruginosa* PAO1– and PAET1-formed biofilms and quantitatively represented between 25 to 35% of the sample, as shown in the pixel color quantification graph (**Figure 1C**). This result demonstrates that both species can survive and coexist in a dual-species biofilm. Apparently, the presence of both morphotypes R and S of *M. abscessus* is higher in the *P. aeruginosa* PAET1 biofilm than in PAO1, consistent with the results obtained in **Figure 1A**, where growth inhibition was higher for the PAET1 strain using the same experimental conditions than for the PAO strain. Regarding the structure of the biofilm, it was observed through orthogonal projections that *P. aeruginosa* and *M. abscessus* were heterogeneously distributed along the biofilm without predominantly occupying any specific area (**Supplementary** Figure 2).

In view of the results obtained, a possible direct inhibitory effect between *P. aeruginosa* PAO1/PAET1 growth and some molecules released by *M. abscessus* R and S, *E. coli* or *B. thuringiensis* was tested. For this purpose, all bacteria were cultivated on agar plates in close streaks where no inhibitory effects were observed on the growth of any bacteria by the presence of the others (**Supplementary** Figure 5A). Kinetic curves of *P. aeruginosa* PAO1 and PAET1 planktonic growth were also generated by cocultivation with culture supernatants of the other bacteria, and similar to the solid agar cultures, no significant inhibitory effect on the growth of *Pseudomonas* was reported (**Supplementary** Figure 5B and 5C).

### P. aeruginosa reduces the development of M. abscessus biofilms

Although infection by *M. abscessus* in CF patients generally occurs in patients previously infected with *P. aeruginosa*, we aimed to study the effect of *P. aeruginosa* on well-developed biofilms of *M. abscessus*. Over a mature *M. abscessus* biofilm (120 h old), the addition of *P. aeruginosa* is directly related to a clear reduction in the development of the *Mycobacterium* biofilm (both for R and S biofilms) (**Figure 2A**), similar to the opposite case (**Figure 1A**). In detail, biofilm inhibition was notably higher in the *M. abscessus* S morphotype than in the R morphotype for all the conditions tested. Additionally, *P. aeruginosa* PAO1 induced a higher growth inhibitory effect in both *M. abscessus* biofilm morphotypes than the *P. aeruginosa* PAET1 strain. These results obtained by GFP quantification were corroborated using confocal microscopy (**Figure 2B**). The images show that after treatment of an *M. abscessus* biofilm with *P. aeruginosa,* both bacteria form a combined biofilm in which *P. aeruginosa* expands rapidly due to its competitive advantage in growth rate. The detection of mycobacteria by GFP in coculture conditions was reduced compared with that of the control biofilms (without *P. aeruginosa*), and a lower presence of *M. abscessus* was also detected in the S morphotype compared to the R in combination with both *P. aeruginosa* PAO1 and PAET1 (**Figure 2B**).

**Figure 2:**
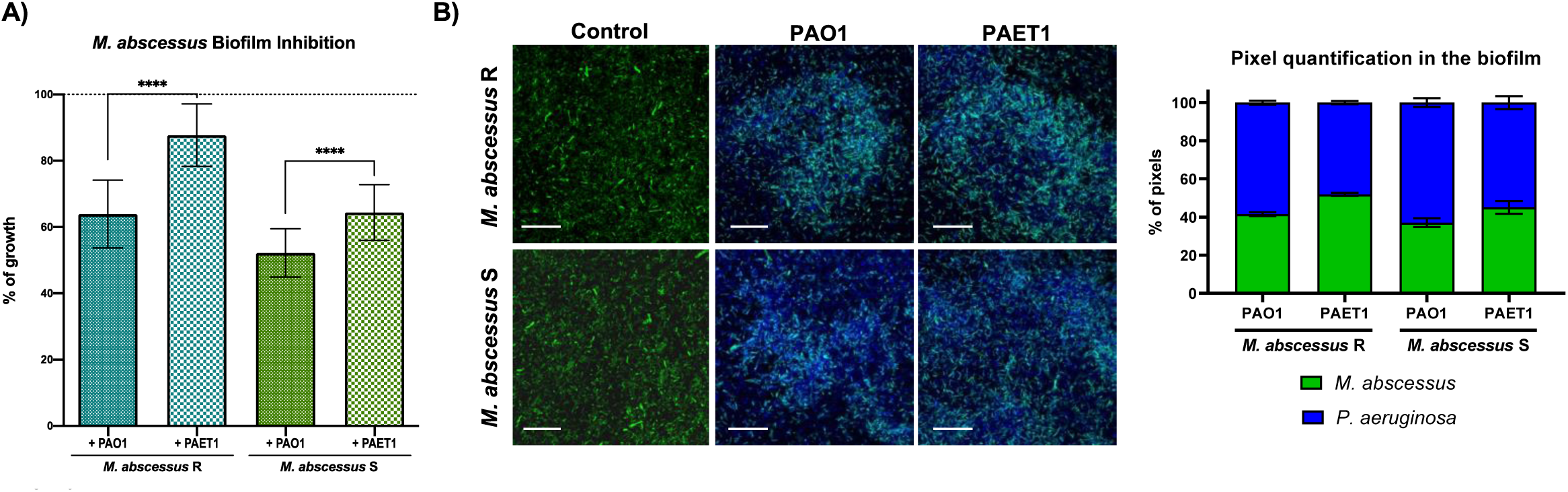
Effect of the *P. aeruginosa* PAO1 and PAET1 strains on *M. abscessus* R and S biofilm development. **A**) Graph represents the inhibition of *M. abscessus* biofilms when cocultured for 72 h with *P. aeruginosa* by measuring GFP expression. The control corresponds to 100% of single *M. abscessus* biofilm growth by analyzing GFP expression (dotted line). Data are presented as the mean ± SD of three independent experiments including six replicates in each experimental condition, and significant differences were reported using the Mann‒Whitney *t* test, *\*\*\*\*, p*<0.0001. **B)** Confocal microscopy representative coculture biofilm images and pixel color quantification. Z-stack compositions of *M. abscessus* variant 120 h biofilms in combination with *P. aeruginosa strains*. DAPI stained all cells blue, and green fluorescent protein (GFP) of plasmid pETS218 allowed mycobacterial cells to be detected in green. Scale bars correspond to 20 µm. Pixel quantifications are represented as the mean values ± SDs from three representative images of each condition.

### Coinfection of M. abscessus and P. aeruginosa in bronchial epithelial cells reduces the viability and immune response triggered with respect to single P. aeruginosa infection

For determination of the effect of bacterial coinfection in the CF context, CFBE41o-(homozygous for the ΔF508 CFTR mutation) and 16HBE14o-(usually differentiated bronchial epithelial cells) bronchial epithelial cells were infected with *P. aeruginosa*, *M. abscessus, B. thuringiensis* or combinations of bacteria. The results showed a reduction in viability of approximately 10-30% in infected cells with respect to uninfected cells in all cases (**Figure 3A**). Remarkably, while single infection with *M. abscessus* variants or *B. thuringiensis* reduced viability up to 5-15% in both cell lines, when cells were coinfected with either of the two *P. aeruginosa* strains, viability was significantly diminished with respect to single infection (Figure 3A). In all cases, the viability levels of coinfected cells were similar to those obtained with single infection of *P. aeruginosa* strains, with the exception of the *P. aeruginosa* strains PAO1 and PAET1, and *M*. *abscessus* R coinfected cells, in which lower viability was obtained (Figure 3A). No differences were obtained between cell lines, indicating that the inhibitory effect of the bacteria is not dependent on the epithelial characteristics.

**Figure 3:**
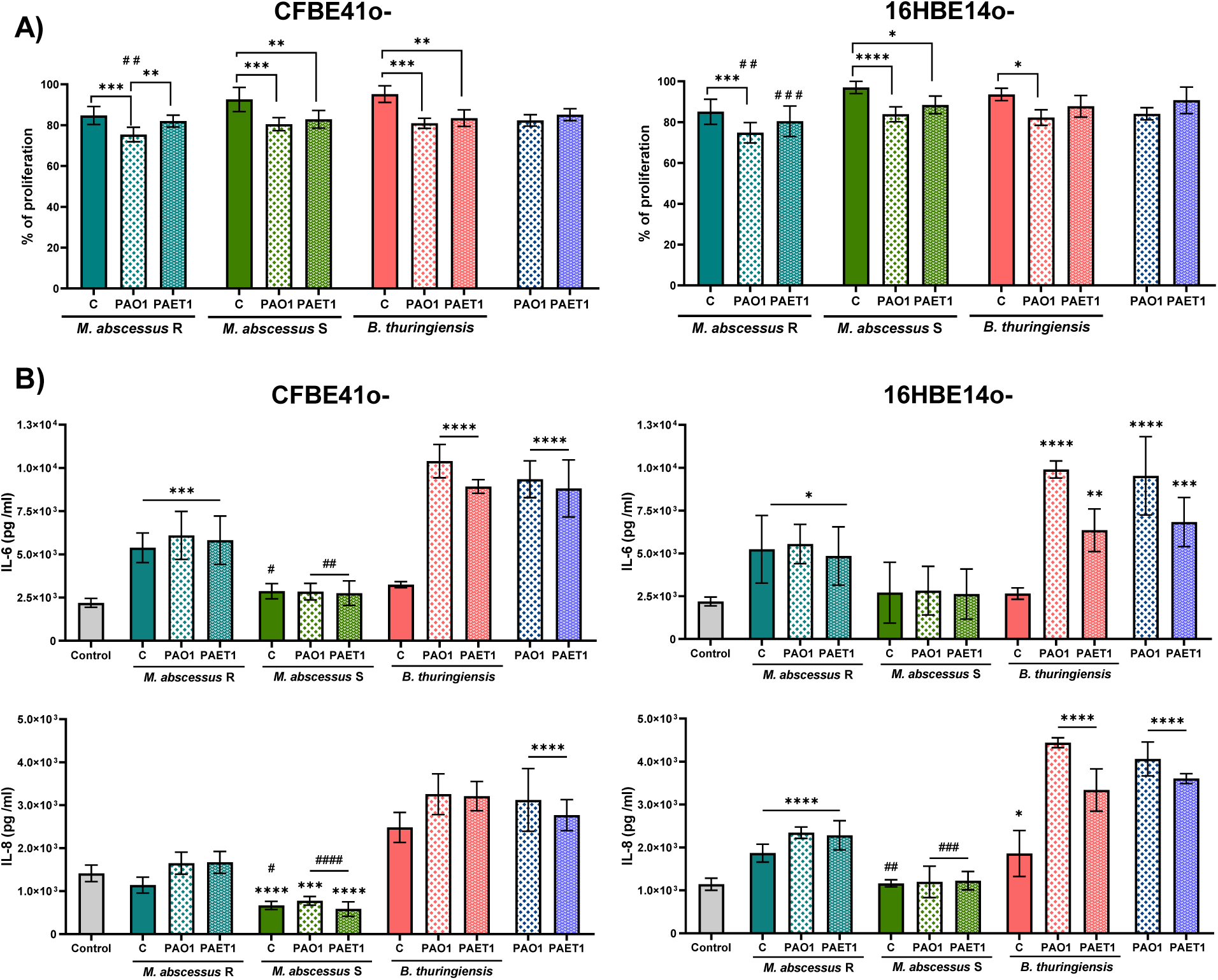
Bronchial epithelial cell viability and proinflammatory cytokine production after *P. aeruginosa* PAO1/PAET1, *M. abscessus* R/S and *B. thuringiensis* infections. **A**) Viability of CFBE41o– and 16HBE14o-cell lines after bacterial infections. Presto blue cell viability assay results in which data are expressed with respect to uninfected wells (considered as 100% cell viability). Data are represented as the mean ± SD from three independent experiments. Significance was analyzed using one-way ANOVA and Tukey’s multiple comparisons test. *, *p*<0.05; *\*\*, p*<0.01; *\*\*\*, p*<0.001; *\*\*\*\*, p*<0.0001. Comparison of cocultures with individual PAO1/PAET1 infections: ^##^, *p*<0.01; *^###^, p*<0.001. B) Cytokine (IL-6 and IL-8) production detected in culture supernatants from bronchial epithelial cell infections. Statistical differences were determined using one-way ANOVA and Dunnett’s test, which compares each column with the control. Data are represented as the mean ± SD from three independent experiments; *, *p*<0.05; *\*\*, p*<0.01; *\*\*\*,* p<0.001; *\*\*\*\*, p*<0.0001. Comparison of results obtained in *M. abscessus* R infections and those in *M. abscessus* S infections were determined using one-way ANOVA, Tukey’s multiple comparisons test; ^#^*, p*<0.05; ^##^, *p*<0.01; *^###^, p*<0.001; *^####^, p*<0.0001.

Differences were observed for each culture condition when IL-6 and IL-8 production was analyzed. When cytokine production was analyzed in cell culture supernatants, *P. aeruginosa*-infected cell lines showed the highest amount, indicating an elevated inflammatory response (**Figure 3B**). Single infections with *M. abscessus* R and S and *B. thuringiensis* showed different stimulatory capabilities, while *M. abscessus* S did not induce the production of cytokines, *B. thuringiensis* triggered only IL-8 production in both cell lines, and *M. abscessus* R triggered IL-6 production in both cell lines and IL-8 only in the control cell line (**Figure 3B**).

The most relevant result was observed in cells coinfected with both bacteria. *M. abscessus* coinfection was able to inhibit the higher production that was reported by *P. aeruginosa* single-infected cells, and the release of IL-6 and IL-8 was reduced to levels similar to those triggered by *M. abscessus* single infection (**Figure 3B**). *M. abscessus* S and *P. aeruginosa* (both strains) coinfection reduced the production of cytokines to levels similar to those in the control cells. Strikingly, this effect observed with *M. abscessus* was not observed when *B. thuringiensis* was coinfecting with *P. aeruginosa*, triggering cytokine levels similar to those produced by *P. aeruginosa* single infections (Figure 3B). No differences were observed between cell lines.

### M. abscessus and P. aeruginosa coinfection accelerates G. mellonella larval death

The *G. mellonella* larvae *in vivo* model presents an immune system broadly similar to mammalian innate immunity being useful for the characterization of microbial infections and host-pathogen interactions. Infection by both *P. aeruginosa* PAO1 and PAET1 strains is lethal for *G. mellonella* larvae. However, *M. abscessus* infections need prolonged times (up to 144 h) and high concentrations (10^6^ cfus/larva) to become lethal (see Materials and Methods section and **Supplementary** Figure 3). Coinfection with *M. abscessus* R and S but not with *B. thuringiensis* accelerated the survival decline in the larvae in both strains and all concentrations tested compared to the infections of *P. aeruginosa* strains alone (**Figure 4**). This overtaking lethality was clearly observed, being significant for all *M. abscesssus* and *P. aeruginosa* coinfection combinations tested and specially in the of PAO1 plus *M. abscessus* R coinfection. We corroborated that the causal agent of larval death was the presence of *P. aeruginosa* PAO1 or PAET1 in the larvae by detecting the luminescence in their interior trough *lux* detection in a luminescence analysis (**Supplementary** Figure 4). As Figure 4 shows, *B. thuringiensis* coinfection did not modify larval survival rates.

**Figure 4:**
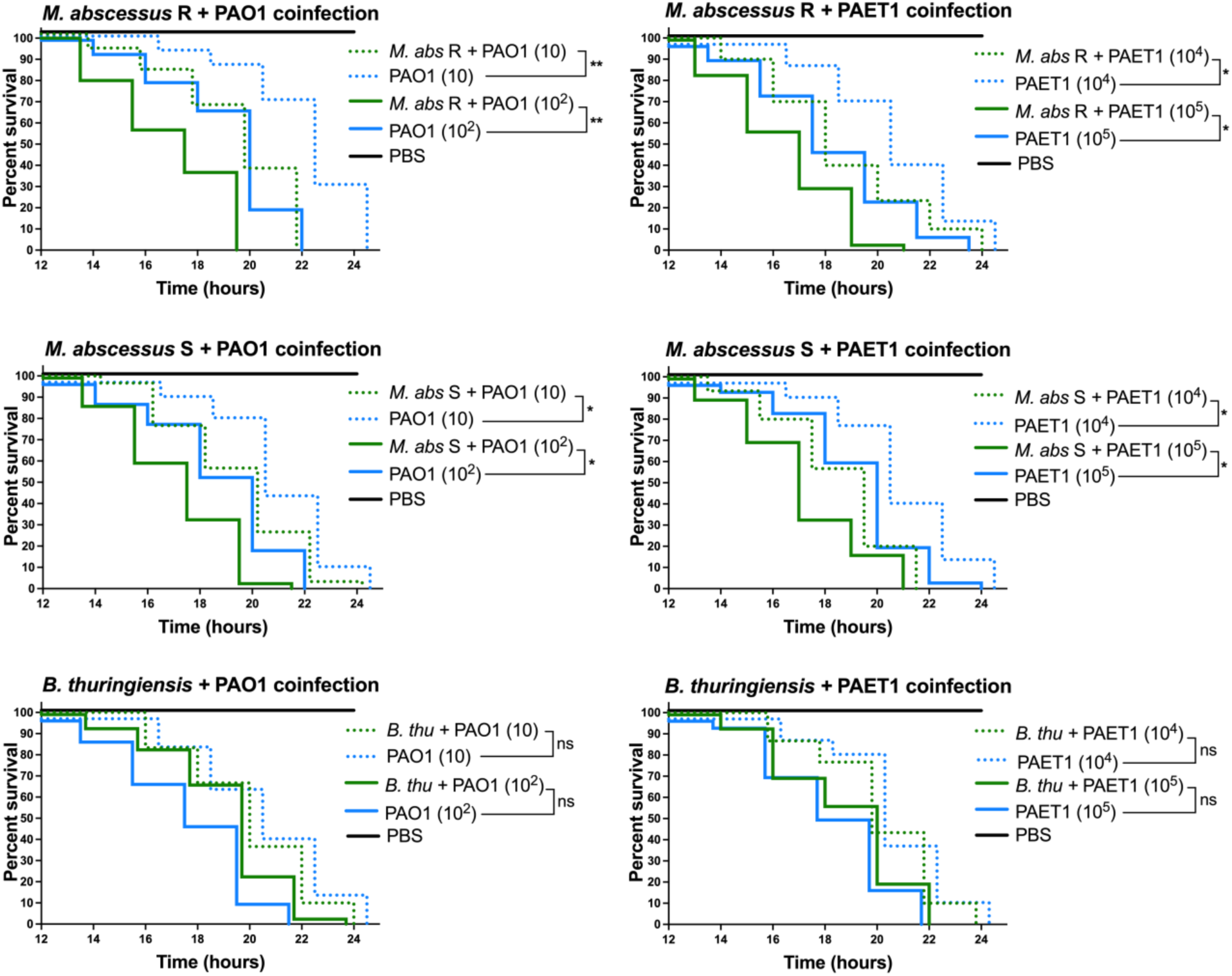
*B. thuringiensis* and *M. abscessus* coinfecting with *P. aeruginosa* in *G. mellonella* Kaplan– Meier survival curves. Thirty larvae (n=30) of each condition were monitored from 12 h post-infection every 2 hours to verify their survival rates when *M. abscessus* (10^4^ CFU/larva) or *B. thuringiensis* (10^5^ CFU/larva) was coinfected with *P. aeruginosa* PAO1 and PAET1 at different concentrations. Data were analyzed by comparing the curves of individual *P. aeruginosa* infections with coinfection of *M. abscessus* plus *P. aeruginosa* at the same concentration using the Mantel‒Cox survival test; *\*, p*<0.05; *\*\*, p*<0.005.

### M. abscessus and P. aeruginosa coinfection restricts the immune response of G. mellonella larvae

*G. mellonella* larvae produce several antimicrobial peptides (AMPs) and enzymes in response to infection. The results of immune-relevant *G. mellonella* gene expression showed a significant decrease in the transcriptional induction for several genes tested, comparing *P. aeruginosa* individual infections with different coinfection conditions (**Figure 5**). For gloverin, cecropin D, transferrin, moricin, lysozyme, insect metalloproteinase inhibitor (IMPI) and nitric oxide synthase (NOS) gene expression, a comparable trend was observed between all the infections assayed. Briefly, high induction of gene expression in PAO1– and PAET1-infected larvae was observed, with lower induction of gene expression in larvae coinfected with *B. thuringiensis* and *P. aeruginosa* strains and notably lower gene expression in the case of larvae infected with both *M. abscessus* R and S alone and coinfected with *P. aeruginosa* (**Figure 5**). These data indicated that the presence of *M. abscessus* generated a decrease in gene expression compared with the infection of single *P. aeruginosa* strains, leaving the induction of larvae coinfected by both bacteria at the levels established by the mycobacteria alone.

**Figure 5:**
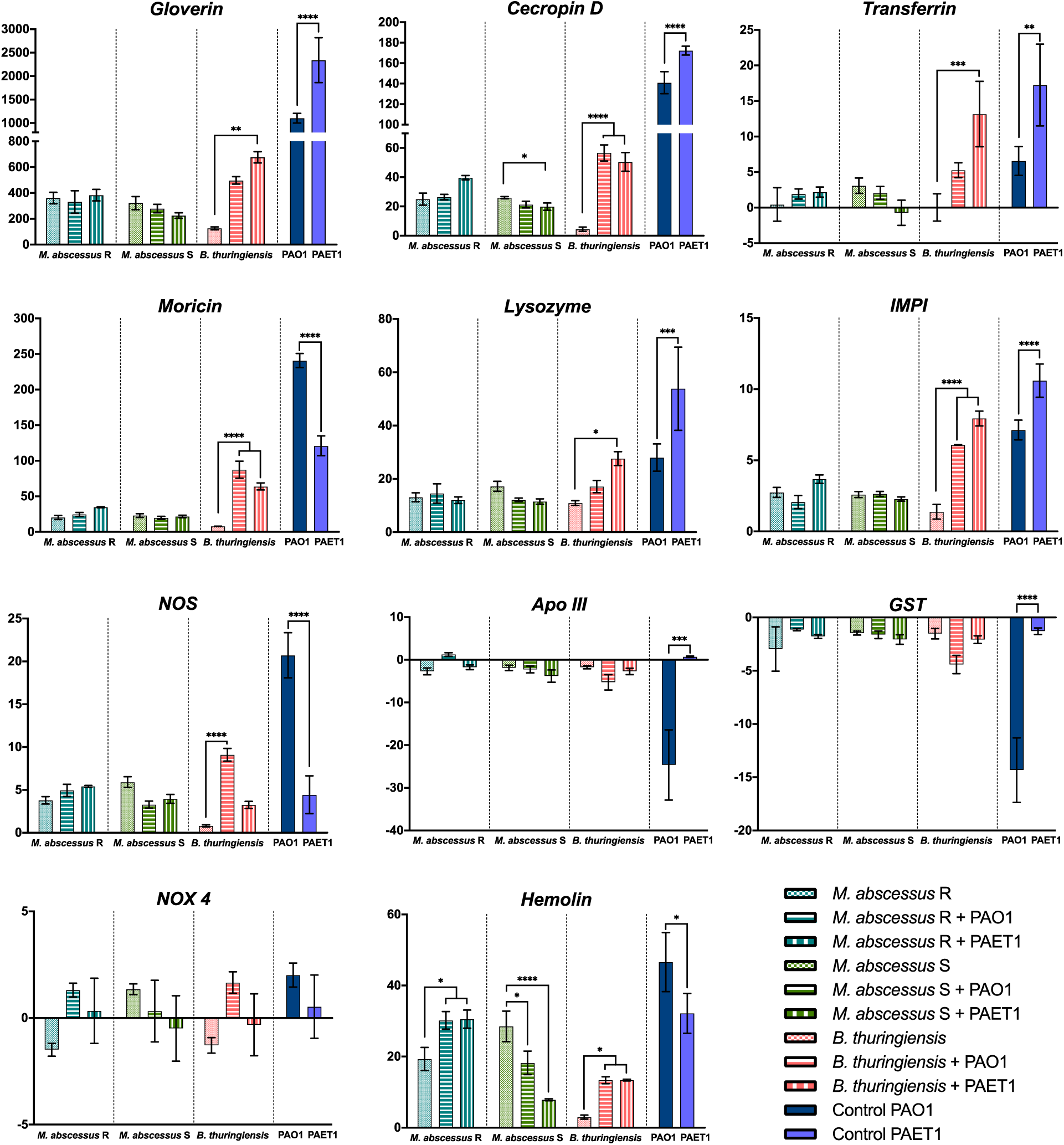
Expression of *G. mellonella* immune-relevant genes under different infection conditions. The fold changes indicate the number of times above or below the gene expression compared with the control (larvae infected with PBS) obtained by RT‒PCR analysis. Data are presented as the mean ± SD of the fold changes for each condition. Significant differences were established using one-way ANOVA. *, *p*<0.05*; **, p*<0.01*; **, p*<0.001*; ****, p*<0.0001*. Apo III*: Apolipophorin III, *IMPI*: insect metalloproteinase inhibitor, *GST*: glutathione S-transferase, *NOX 4*: NADPH oxidase, *NOS*: nitric oxide synthase.

Apolipophorin III (Apo III) and glutathione S-transferase (GST) resulted in a similar trend, with no significant differences detected compared to the control (PBS-infected larvae) in any conditions except for the *P. aeruginosa* PAO1-infected larvae, which showed evident repression of both genes. NADPH oxidase (NOX4) gene expression was the only one that did not show significant differences in any of the infections tested. In the case of hemolin, increased gene expression was observed in *M. abscessus*-infected larvae, much higher than in the other studied genes and higher than in *B. thuringiensis* infection conditions. This result contradicts the rest of the immuno-relevant gene tendencies. Hemolin acts as an opsonin and mediates the interaction between microorganisms and hemocytes, stimulating phagocytosis. In this work, we have also shown that *M. abscessus* R and S could be phagocytosed by *G. mellonella* hemocytes (**Supplementary** Figure 6A) and progressively eliminated from the larva (**Supplementary** Figure 6B), which would explain this contradictory result compared to the rest of the genes.

## Discussion

Infections of the respiratory tract epithelium are the leading cause of morbidity and mortality in CF patients. *P. aeruginosa* is the most relevant pathogen due to its robust virulence systems and biofilm-forming capacity, which trigger increased tolerance to antibiotics and resist phagocytosis (9, 23). Although *P. aeruginosa* is the most common and prevalent pathogen in CF patients, other microorganisms, such as *S. aureus*, *Haemophilus influenzae*, *Stenotrophomonas maltophilia*, NTM, and *Burkholderia* species, are presently forming a polymicrobial environment. Within the NTM group, *M. abscessus* is the most relevant pathogen in CF patients since its multidrug-resistant and biofilm formation capabilities (20, 24) widely increase antibiotic resistance and prevalence. In this work, the relationship between *P. aeruginosa* PAO1 (laboratory strain)/PAET1 (clinical isolate) and *M. abscessus* R/S morphotypes was studied for the first time in different contexts of critical interest: biofilm development, bronchial epithelial cell *in vitro* cultures and *G. mellonella* larvae *in vivo* infection.

Regarding biofilm development, the results indicate a decrease in the progression of *P. aeruginosa* biofilms when live *M. abscessus* R and S were introduced, independent of whether the biofilm was already well established (mature biofilm) or in coculture biofilm growth. Inhibition of *Pseudomonas* biofilm development by cocultivation with other pathogens commonly found in the lungs of CF patients, such as *Aspergillus,* or potential probiotics, such as lactobacilli, has already been reported (25, 26). Nevertheless, no inhibitory effect has been reported previously for mycobacteria. This inhibition could be related to the capacity of *M. abscessus* to inactivate quorum sensing quinolone signals from *P. aeruginosa* (27). In addition, the results showed that both species coexist in dual-species biofilms, with *P. aeruginosa* being the dominant species, most likely related to its competitive advantage in terms of growth since *M. abscessus* has a division time of 4 to 5 h, while *P. aeruginosa* has a division time of only 25 to 35 min (28, 29). Conversely, biofilms of *M. abscessus* are also inhibited by the presence of *P. aeruginosa,* which outcompetes mycobacteria, as previously reported in biofilm coculture experiments (30). The mechanism responsible of this inhibition remain still unknown and *P. aeruginosa* interbacterial competition secretion systems and quorum sensing seem to be not involved (31). Distinctly, the growth of S morphotype biofilms, despite forming biofilm more easily by the abundance of GPLs in its cell wall (32), is inhibited more by the presence of *P. aeruginosa* than R morphotype biofilms, which could be related to the aggregative phenotype that confers higher resistance to antibiotics as well as structural rigidity (33). These results indicate a decrease in both species biofilm progression when coculturing. It can be hypothesized that this effect could not be due to direct inhibition but rather a consequence of the combined growth of both species sharing the same space and competing for the same culture medium. This hypothesis is partly supported by the lack of direct inhibition observed in agar plate cultures using *M. abscessus* R and S cultured close to *P. aeruginosa* PAO1 and PAET1 or in planktonic growth kinetics curves of *P. aeruginosa* PAO1 and PAET1 in combination with culture supernatants of the rest of bacteria, in which no significant effect on the growth of *P. aeruginosa* was highlighted (**see Supplementary** Figure 5). Also, the limitation in the progression of the *P. aeruginosa* PAO1 biofilm was observed in co-culture with a non-pathogenic environmental mycobacteria such as *Mycobacterium brumae* (data not shown). It has been demonstrated that in dual-species biofilms, antibiotic treatment to eliminate *P. aeruginosa* supposes a competitive advantage for *M. abscessus* development (17). In any case, *M. abscessus* establishes itself effectively and, despite its difference in growth rate, prevails in dual-species biofilms, demonstrating its resistance and survival capabilities (34).

Regarding the effect of coinfection on bronchial epithelial cell viability *in vitro*, it has been shown that coinfection results in higher mortality compared to individual *M. abscessus* and *B. thuringiensis* infections, as expected. However, as previously reported (35), the viability of the cell lines tested, CFBE41o– and 16HBE14o-, was not excessively compromised, being approximately 80% of the control in all tested conditions. *M. abscessus* and *P. aeruginosa* coinfections correlate with an elevated pulmonary function decline and worse disease severity (36, 37); therefore, lower viability of the epithelial cells could be involved in these complications. The inflammatory response triggered by epithelial cells shows a clearly exacerbated production of proinflammatory cytokines, IL-6 and IL-8, in response to PAO1 and PAET1 single infections (see Figure 3). This immune response against *P. aeruginosa* is harmful to patients, as excessive inflammation is responsible for most of the morbidity and mortality in CF (38–40). Surprisingly, coinfection of *M. abscessus* and *P. aeruginosa* with epithelial cells showed an evident inhibition of the proinflammatory response promoted by the mycobacteria since single *M. abscessus*-infected cells presented similar production levels as the coinfected cells, being especially prominent in the case of *M. abscessus* S. In general, microorganisms that occupy the airways together with *P. aeruginosa* also induce a proinflammatory response, as in the case of *B. cenocepacia, H. influenzae, S. aureus* and *Candida spp.* among others (41, 42). A comparable inflammatory reduction effect to that observed for *M. abscessus* in our results has only been demonstrated in *P. aeruginosa* coinfections with *Prevotella spp*. an anaerobic gram-negative bacterium belonging to the respiratory tract microbiota (43). In the case of *M. abscessus*, previous works demonstrated that bronchial epithelial cells generate a hyporesponsive response since *M. abscessus* S fail to elicit a Toll-like receptor 2 (TLR2) response and only a precise cell fraction from R morphotype exhibits TLR2 stimulating activity (44), which is necessary for the activation of the signaling cascade that triggers the secretion of the cytokines IL-6 and IL-8 (45). This lack of immune response activation observed is particularly high in the case of the S morphotype (as supported by our results) since the presence of GPL in its cell wall masks other molecules that would activate TLR2 signaling (46). In most cases, this factor also explains why initial *M. abscessus* colonization of lung airways occurs with an S morphotype strain. In the CF context, a beneficial immune response consists of the capacity to activate an inflammatory response to manage infection but to avoid an exuberant response that can be harmful (47). In the present study, cocultures of *M. abscessus* and *P. aeruginosa* showed less inflammatory activation and the highest loss of viability among infected cells, while cocultures of *B. thuringiensis* and *P. aeruginosa* showed a high inflammatory response, leading to a high loss of viability. These results could indicate that an excessive proinflammatory response and significant immune inhibition could be equally harmful to infected lung epithelial cells.

The results of the *in vitro* cultures were corroborated by infections in the *G. mellonella* larva model, in which an increased lethality was observed in coinfection conditions with respect to the single *P. aeruginosa* infection. *G. mellonella* is highly susceptible to *P. aeruginosa* infections since 100% larval lethality is reached in less than 24 h with few inoculated bacteria (10-30 CFU) (48, 49) although there are important variations in pathogenicity and lethality between strains commonly used in research (PAO1 in this study) and clinical isolates (PAET1 in this study) (35, 50). However, *M. abscessus,* despite being pathogenic for larvae, requires much higher doses and prolonged infection times than *P. aeruginosa* to be equally lethal ((51, 52), and **Supplementary** Figure 3). In addition, in agreement with our *in vitro* results, the immune response of *G. mellonella* larvae was clearly inhibited under coinfection conditions (Figure 5). The gene expression of most proteins and enzymes related to larval immunity showed a significant decrease compared with that of *P. aeruginosa* strain single infections (53). In detail, relevant AMPs involved in antimicrobial activity, such as gloverin, cecropin D and moricin, were clearly inhibited in coinfection conditions. All of them act against the bacterial cell wall by interacting with LPS and other components, especially affecting the outer membrane of gram-negative bacteria (54–56); specifically, moricin is capable of forming pores in gram-positive and gram-negative cell walls (57). Under coinfection conditions, insect metalloproteinase inhibitor (IMPI) expression was also inhibited; this peptide has only been described thus far in *G. mellonella* (58) and inhibits the activity of proteases secreted by invading microorganisms (59). In this sense, it is well known that *P. aeruginosa* produces various proteases as virulence factors (60); thus, high levels of IMPI would potentially slow the progression of *P. aeruginosa* infection. Furthermore, IMPI inhibits the activation of phenoloxidase, the activator enzyme of the melanization cascade (61). The produced melanin is deposited around the pathogenic microorganism, limiting its ability to damage the larva, but in parallel to melanin production, dangerous substances and reactive substances are also produced, resulting in tissue damage and cell death, which could be self-defeating for the host (62). Additionally, the expression levels of immune-relevant proteins and enzymes, transferrin, lysozyme and nitric oxide synthetase (NOS), are notably inhibited when *M. abscessus* and *P. aeruginosa* are infecting simultaneously. Lysozyme is a cationic peptide with muramidase activity (63) that degrades peptidoglycan, making it more effective against gram-positive bacteria and, similar to IMPI, preventing an excessive melanization response (64). Transferrin mediates nutritional immunity by sequestering iron from invading pathogens, limiting their ability to grow and develop (65). *P. aeruginosa* needs substantial amounts of iron to grow and produces two siderophores to take up this metal (66), so high levels of transferrin in the hemolymph are beneficial to host survival. The production of nitric oxide by NOS under infection conditions is increased, although its role in immunity is not entirely clear (67), although it is hypothesized that it is transformed into reactive nitrogen species that are highly toxic to bacteria but equally toxic to host cells (68). Moreover, the expression levels of glutathione S-transferase (GST) and apolipophorin III (Apo III) are clearly repressed only in *P. aeruginosa* PAO1-infected larvae, while they remain unchanged in the other conditions, including *P. aeruginosa* PAET1, which indicates differences in the immune response against both *P. aeruginosa* strains. GST is a detoxification enzyme that protects the host from oxidative stress (69). To reduce pathogen progression, larvae induce a prophenoloxidase response, which triggers the production of reactive oxygen species to damage bacteria, although it also affects host cells (70); therefore, it would make sense to inhibit antioxidative enzymes, such as GST, under severe infections. Apo III is a lipid transport protein with relevant immune effects since it mediates hemocyte adhesion, phagocytosis and nodule formation (71). *P. aeruginosa* serine protease IV was shown to degrade Apo III (72, 73); however, the results of our study show that Apo III levels are also reduced at the transcriptional level. In addition to the protease activity of *P. aeruginosa*, low levels of Apo III in hemolymph could be reinforced by a repressive effect on its synthesis. NADPH oxidase (NOX4) is the only enzyme that did not present significant differences between infected and uninfected larvae. This enzyme produces superoxide and reactive oxygen species in response to infection, suggesting that it is not overexpressed under the infection conditions tested in this study due to the production of these toxic agents by the prophenoloxidase pathway (74), which is highly induced, as previously mentioned.

The repressive effect of *M. abscessus* R and S, together with *P. aeruginosa* PAO1 and PAET1 coinfections, was observed in most of the genes evaluated. However, as the only exception, an induction of transcriptional expression of the hemolin gene in *M. abscessus*-infected larvae (especially the R variant) was shown. Hemolin is an immunoglobulin-like protein with no direct antibacterial properties associated with hemocytes in the pathogen recognition process. This protein functions as an opsonin that mediates between bacteria and hemocytes, stimulating the phagocytosis process. Surprisingly, this study shows the phagocytic capacity of *G. mellonella* hemocytes against *M. abscessus* bacilli (**Supplementary** Figure 6), which could be directly related to this singular overexpression detected in mycobacterium-infected larvae.

Overall, the transcriptional expression levels of the different genes investigated show high induction levels in the single infections of *P. aeruginosa*, although slight differences between PAO1 and PAET1 strains are marked, showing different responses. In *M. abscessus*-infected larvae, expression levels are notably lower and remain at similar levels in both single and coinfections. This repression of the larvae innate immune response could be attributed to the extensive phagocytosis of mycobacteria by hemocytes, which may proliferate and thereby limit the antibacterial activity of these cells, consequently hindering a robust specific response against *P. aeruginosa*. Additionally, prior studies have highlighted *M. abscessus* specific affinity for colonizing and infecting the fat body tissue of *G. mellonella* larvae (75), which serves as the primary source of antimicrobial peptides, enzymes, and antibacterial proteins secreted into the hemolymph (59). Hence, the presence of *M. abscessus* in the fat body could account for the diminished production of these molecules that correlates with the acceleration in lethality in the coinfection conditions. Notably, larvae infected with *B. thuringiensis* follow trends similar to those of individual *P. aeruginosa* infections, although interestingly showing lower levels than these.

The observed results in the immune response triggered by bronchial epithelial cells and *G. mellonella* larvae are compatible with the decreased response degree under coinfection and contrary to the overexpressed response under *P. aeruginosa* single infections. The hyperinflammatory response produced in the airways by *P. aeruginosa* infection is well described in patients (76), producing more severe respiratory malfunction usually treated with corticosteroids (77). In the case of *M. abscessus*, it has been shown in a murine model of CF that the proinflammatory response produced is higher in the R morphotype than in the S morphotype (78), in accordance with the IL-6 and IL-8 differences detected in our study. The immune repression shown in the coinfections of *M. abscessus* and *P. aeruginosa* is described for the first time in this work. A disproportionate immune response is dangerous for the host since excess inflammation and the production of toxic compounds for bacteria and the host trigger a decline in survival. However, the weak immune response observed in coinfections with *M. abscessus* is even worse since the bacteria can spread unrestrictedly, triggering lethality at shorter times, as shown in the results of the *in vivo* survival and *in vitro* viability assays in this work. Taking this study as a starting point, and in order to generalize the results, it will be necessary to explore whether the results obtained are replicable using other strains of *P. aeruginosa* and *M. abscessus* beyond those employed in this study, particularly with clinical relevance, which allow establishing a clear pattern of interaction between both species when they are coinfecting simultaneously.

## Materials and Methods

### Bacterial strains and culture conditions

Two different *P. aeruginosa* strains were used: PAO1 (ATCC 15692) as a reference laboratory strain and PAET1, a clinical isolate from a chronic CF-infected patient (79). Both strains were transformed with a plasmid (pETS130lux) expressing a constitutive bacterial luciferase gene cassette (*lux*) as a bioluminescent bioreporter system (80). Both *P. aeruginosa* strains were grown in Luria-Bertani (LB, Sharlab, Barcelona, Spain) agar with 50 μg/ml gentamycin (Gm) for PAO1 and 300 μg/ml carbenicillin (Cb) for PAET1 as a selection marker. For biofilm formation assays, *P. aeruginosa* was grown in tryptic soy broth (TSB, Sharlab) without antibiotics. *Bacillus thuringiensis* (ATCC 10792) was cultured in tryptic soy agar (TSA, Sharlab) or TSB, and *Escherichia coli* MG1655 (ATCC 700926) was grown in LB (agar and broth). The *M. abscessus* DSMZ 44196 (ATCC 19977) original smooth (S) morphotype and the rough (R) variant, a natural mutant of the wild-type strain previously obtained (81), were grown on TSB with agitation and 0.5 mm diameter glass beads (DDBiolab, Barcelona, Spain) to prevent cell aggregation and on TSA agar for colony forming unit (CFU) counting. As previously reported, both variants of *M. abscessus* were transformed with the plasmid pETS218 (82). For further detail regarding the strains used, see **Supplementary Table S1**.

### DNA manipulation and plasmid construction

Molecular biology enzymes and kits were purchased from Thermo Fisher Scientific (Madrid, Spain) and used according to the manufacturer’s instructions. DNA amplification was performed by PCR using DreamTaq MasterMix (2X) or High-Fidelity PCR Enzyme Mix (Thermo Fisher Scientific). The primers used are listed in **Supplementary Table S2**. Plasmid pETS218 was constructed by cloning the promoter region of the class Ib ribonucleotide reductase (encoded by the *nrdHIE* genes) from *M. brumae* (GenBank assembly accession: GCA_900073015.1). All other recombinant DNA manipulations were performed using standard procedures (83). For construction of mycobacterial *nrdH* transcriptional GFP fusions, a 537 bp long fragment encompassing the *nrdH* promoter region was amplified by PCR using the primer pair MbruPnrdHIE-For-MbruPnrdHIE-Rev and *M. brumae* genomic DNA; the obtained DNA fragment was cloned into pJET1.2 and transformed into *E. coli* DH5α cells. *BamHI* and *Apa*I restriction enzymes were used for fragment digestion and cloning into pFPV27 (84) to generate pETS218. The absence of mutations introduced during cloning was verified via DNA sequencing. *M. abscessus* S and R morphotypes were grown in TSA plus 50 μg/ml kanamycin (Kn) when they were transformed with pETS218 as we previously described (82).

### Biofilm progression inhibition quantification

Biofilm quantifications were performed using a microtiter plate screening assay previously optimized, validated and published in our group (85). Briefly, mature *P. aeruginosa* PAO1 and PAET1 biofilms (72 h growth) were cocultured with *B. thuringiensis*, *E. coli* and *M. abscessus* R and S for 24h further. For this purpose, ON cultures of *B. thuringiensis* and *E. coli* were adjusted to an OD_550_ nm of 1 and appropriately diluted to obtain the desired concentrations. *M. abscessus* R and S were grown for 3 days on TSB cultures, and the concentrations of both live and heat-killed (121 °C for 21 min) cell suspensions were determined by comparison to the MacFarland 1 (McF) turbidity standard. Bacterial culture supernatants were also obtained by centrifugation of these cultures at 12,000 g. Serial dilutions of the bacterial suspensions were plated to verify the accurate concentration in each experiment for all bacteria tested. The level of lux expression (*P. aeruginosa* cells) in the biofilms was quantified using a Spark® multimode microplate reader (Tecan, Männedorf, Switzerland). For simultaneous dual-species biofilm formation experiments, *P. aeruginosa* PAO1 and PAET1 suspensions were mixed with the rest of bacteria or supernatants and grown for 72 h.

### M. abscessus biofilm formation and P. aeruginosa coexistence

Three-day-old liquid cultures of *M. abscessus* R and S transformed with pETS218 plasmid were adjusted to a concentration of 5×10^6^ cfu/ml and grown in TSB medium + 0.2% glucose in Costar® 96-Well Black Polystyrene Plates (200 μl/well) (Corning) at 37 °C in saturation humidity conditions to allow mycobacterial biofilm formation. After 120 h of incubation, 3 washes with PBS were carried out, and suspensions of *P. aeruginosa* PAO1 and PAET1 cultures were added at an OD_550 nm_= 0.1 for an additional 72 h. Finally, the biofilms were homogenized in PBS and quantified by measuring the GFP and *lux* expression associated with *M. abscessus* and PAO1/PAET1 growth, respectively, using a Spark® multimode microplate reader.

### Biofilm analysis through confocal microscopy

The *P. aeruginosa* and mycobacterial biofilms developed in 96-well plates were collected, and bacteria were stained with 4’,6-diamidino-2-phenylindole (DAPI). *M. abscessus* R and S (transformed with pETS218) bacilli were specifically detected by measuring their GFP expression. Representative images were obtained in a Zeiss LSM 800 confocal laser scanning microscope (CSLM, Zeiss, Oberkochen, Germany) using the 63×/1.4 oil objective with excitation wavelengths of 405 and 488 nm for DAPI and GFP, respectively. Five representative Z-stack images were selected and analyzed for pixel color quantification using the color histogram tool from Fiji-ImageJ software and Zen software (Zeiss).

### Galleria mellonella maintenance and infection

*G. mellonella* larvae were fed *at libitum* with an artificial diet as previously described (80). The larvae were kept at 34 °C in a dark environment until they reached their optimum size (approximately 200 mg). *G. mellonella* larvae were injected with 10 µl of bacterial suspension containing the desired concentration through the top right proleg using a 25 µl Hamilton microsyringe (Hamilton, Reno, USA). In previous experiments, lethal and innocuous doses of each bacterium were determined (**Supplementary** Figure 3). Based on these results, the highest safe doses were selected: 10^4^ CFU/larva for *M. abscessus* R and S and 10^5^ CFU/larva for *B. thuringiensis* and *E. coli*. In the case of *P. aeruginosa*, even the lowest dose tested (10 CFU/larva) was lethal. The virulence of the *P. aeruginosa* PAO1 strain was much higher than that of PAET1; therefore, the PAET1 strain dose was adjusted to 10^4^ CFU/larva, similar to 10 CFU/larva of PAO1. Thirty larvae were infected with each condition, *E. coli*, *B. thuringiensis* and *M. abscessus* R/S, and were then injected with the *P. aeruginosa* PAO1 or PAET1 strains. Additionally, larvae infected with only each *P. aeruginosa* strain were used as controls for comparison purposes. Infected larvae were kept at 37 °C to allow the development of infection for 14 h. At that time, five larvae were used to extract RNA, and thirty were monitored for survival for 24 h.

For analysis of the phagocytosis of *M. abscessus* R and S by *G. mellonella* hemocytes and the concentration of mycobacteria in the hemolymph (**Supplementary** Figure 6), the larvae were infected with 10^4^ CFU/larvae of each variant. For collection of hemolymph, larvae were placed on ice for 10 min. Once anesthetized, the anal proleg was cut with a surgical blade, and the hemolymph was collected in a 1.5 ml Eppendorf tube. Hemolymph from 10 larvae was pooled and appropriately diluted and cultured on TSA plates for mycobacterial CFU counts. The hemolymph from the other 10 larvae was centrifuged and washed 3 times with PBS (5 min at 500 g at 4 °C). Precipitated hemocytes were resuspended in PBS and stained with DAPI (blue) (4′,6-diamidino-2-phenylindole) to observe nuclei and with FM 4-64 (red) (N-3-triethylammoniumpropyl-4-6-4-diethylamino phenyl hexatrienyl pyridinium dibromide) for dying hemocyte vesicles and plasma membrane and observed under a confocal microscope (Zeiss LSM 800 confocal laser scanning microscope (CSLM)).

### G. mellonella RNA extraction and immune-relevant gene expression analysis

At 14 h post-infection, five larvae were placed on ice for 10 min. The larvae were homogenized using a rotor-stator homogenizer (IKA, Staufen, Germany), and 30 mg of the homogenate was used for RNA extraction following the GeneJET RNA Purification Kit instructions (Thermo Fisher Scientific). RNA obtained was treated with 10x TURBO DNase (Life Technologies, Carlsbad, USA) for 1 h to eliminate possible DNA contamination. DNA absence was verified by PCR amplification of the 18S rRNA housekeeping gene using genomic DNA of *G. mellonella* as a positive control. RNA was quantified in an M200 PRO microplate reader (Tecan). RNA to cDNA reverse-transcription was performed using Maxima Reverse Transcriptase (Thermo Fisher Scientific) with Oligo (dT)18 primers (Thermo Fisher Scientific) by following the manufacturer’s instructions. The cDNA obtained was stored at –20 °C until use. Quantitative real-time PCR (qRT-PCR) was performed using PowerUp™ SYBR™ Green Master Mix (Applied Biosystems, Foster City, CA, USA) in a StepOnePlus™ Real-Time PCR System (Applied Biosystems) according to the manufacturer’s protocol. All qRT-PCRs used specific primers of *G. mellonella* immune-relevant genes listed in **Supplementary Table 2**.

### Bronchial epithelial cell line culture and infection

Two human bronchial epithelial cell lines were used: CFBE41o-, isolated from a CF patient homozygous for the ΔF508 CFTR mutation, and 16HBE14o-, isolated from a cardiopulmonary patient with overexpression of CFTR (35). Cells were maintained in Dulbecco’s modified Eagle’s medium: nutrient mixture F12 (DMEM/F12; Thermo Fisher Scientific) with 10% (v/v) decomplemented fetal bovine serum (dFBS) (Gibco, Paisley, UK) and 1% (v/v) penicillin‒streptomycin (Thermo Fisher Scientific). Cells were maintained in a humidified incubator at 37 °C and 5% (v/v) CO_2_ (ICOmed, Memmert). For infection experiments, cells (5×10^4^ cell/well) were incubated in 96-well plates (Corning) using antibiotic– and serum-free medium for 3 h to allow cell adhesion. Subsequently, the cells were infected at a multiplicity of infection (MOI) of 50:1 for *P. aeruginosa* PAO1 and PAET1 (50:1) and of 10:1 for *M. abscessus* R / S and *B. thuringiensis*. Infections with each bacterium as well as coinfections combining both strains of *P. aeruginosa* and *M. abscessus* were carried out. Three hours later, the wells were washed 3 times with PBS to remove extracellular bacteria, and the plates were incubated for 24 hours. The supernatant was then removed and stored at –80 °C for cytokine measurements. The effect of *P. aeruginosa*, *M. abscessus* and their combination on cell viability was analyzed by Presto Blue assay (Thermo Fisher Scientific) following the manufacturer’s instructions, using uninfected cells as a control. Cell infections were performed in triplicate in three independent experiments.

### IL-6 and IL-8 detection by enzyme immunoassays (ELISAs) in cell supernatants

IL-6 and IL-8 production was measured in cell culture supernatants using ELISA kits according to the manufacturer’s instructions (Becton and Dickinson, BD, San Diego, USA). Briefly, wells were coated with IL-6 or IL-8 capture antibody diluted in coating buffer (0.1 M sodium carbonate, pH 9.5) overnight at 4 °C. Plates were then washed and blocked with assay diluent (PBS with 10% FBS) for 1 h at room temperature (RT). Plates were prepared with standards diluted in assay diluent, and samples were incubated for 2 h at RT. After washing, streptavidin-horseradish peroxidase conjugate mixed with biotinylated detection antibody was then added to each well for 1 h. Finally, tetramethylbenzidine (TMB) substrate solution and hydrogen peroxide (BD, San Diego, USA) were added to the plates for 30 min in the dark. Absorbance (OD_630 nm_) was measured in an Infinite 200 PRO microplate reader (Tecan) and analyzed with Magellan software (Tecan).

## Author Contribution

The manuscript was written through the contributions of all authors. Víctor Campo performed the experiments. Esther Julián and Eduard Torrents directed the research, revised the experimental data, and wrote the manuscript. All authors have approved the final version of the manuscript.

## Acknowledgments

This study was supported by grants from the Ministerio de Ciencia, Innovación y Universidades and Agencia Estatal de Investigación (AEI), Spain, cofunded by Fondo Europeo de Desarrollo Regional, FEDER, European Union (RTI2018-098777-B-I00, PID2021-122331OB-I00, PID2021-125801OB-100, PDC2022-133577-I00 and PLEC2022-009356), the CERCA programme and *AGAUR-Generalitat de Catalunya* (2021SGR-0092 and 2021SGR-0145), the European Regional Development Fund (FEDER), the Catalan Cystic Fibrosis association and Obra Social “La Caixa”. VC-P is a recipient of a Ph.D. contract (FI) from the Generalitat de Catalunya.

## Data Availability

The datasets used and/or analysed during the current study available from the corresponding author on reasonable request.

