## Supplementary material for "Interplay of *Mycobacterium abscessus* and *Pseudomonas aeruginosa* in coinfection: Biofilm Dynamics and Host Immune Response": Supplemetary information

**Supplementary information**

**Supplementary Table 1: Strains and plasmids.** Detail of the strains and plasmids used in this study, *P. aeruginosa* strains PAO1 and PAET1 were transformed with the pETS130Lux plasmid to quantify the expression of luciferase (*Lux*). The *M. abscessus* strains were transformed with the pETS218 plasmid for constitutive expression of green fluorescent protein (GFP).

| Strains | Observations |
| --- | --- |
| *Pseudomonas aeruginosa* PAO1 | Reference strain for laboratory studies. Compatible with a cystic fibrosis acute infection. |
| *Pseudomonas aeruginosa* PAET1 | Cystic fibrosis strain isolated from a chronic patient (79) |
| *Mycobacterium abscessus* Smooth morphotype | Type strain. (DSMZ 44916) |
| *Mycobacterium abscessus* Rough morphotype | Strain obtained by several passages of the original type strain (81) |
| *Escherichia coli* str. K-12 substr. MG1655 | Model strain in molecular biology. Safe strain used as control. |
| *Bacillus thuringiensis* | Type strain. (CECT 197) |
| *Escherichia coli* DH5α | Versatile strain used for general cloning applications. In the study it was used to preserve the different plasmids: pFPV27, pJET 1.2 + PnrdH, pETS218. All were transformed by thermal shook. |

| Plasmids | Observations |
| --- | --- |
| pFPV27 | Mycobacteria expression, it has a kanamycin resistance gene and another that codes for GFP. |
| pJET1.2 | Positive selection cloning vector |
| pETS130Lux | Plasmid previously constructed to obtain *Lux* constitutive expression in *P. aeruginosa* |
| pETS218 (MbruPnrdHIE + pFPV27) | Plasmid constructed to obtain GFP constitutive expression in *M. abscessus* |

**Supplementary Table 2: List of primers used for pETS218 construction and the RT-PCRs experiments.** The list includes the genes with relevance in the immune response of the *Galleria mellonella* larva against infection by bacteria used in this study.

| Primer name | Sequence 5´-3´ | Observations |
| --- | --- | --- |
| MbruPnrdHIE For | AAGGATCCGCGGTTCGCGACGCCGTC | Primers used to amplify the promoter region of the class Ib ribonucleotide reductase. This sequence was then used to clone into pFPV27 to obtain a constitutive expression of GFP (pETS218) |
| MbruPnrdHIE Rev | AAGGGCCCAGCGCCTTGTAGGTCGCGTT |  |
| 18S rRNA For | ATGGTTGCAAAGCTGAAACT | Housekeeping gene: Used to normalize gene expression data in RT-PCRs. |
| 18S rRNA Rev | TCCCGTGTTGAGTCAAATTA |  |
| Apo III For | AGACTTGCACGCCATCAAGA | Apolipophorin III: activity as pathogen recognition receptor, stimulating the activity of defense peptides, and possessing antimicrobial activity itself. |
| Apo III Rev | TGCATGCTGTTTGTCACTGC |  |
| Gloverin For | AGATGCACGGTCCTACAG | Interacts with lipopolysaccharides (LPS) and inhibit the formation of Gram-negative bacterial outer membrane. |
| Gloverin Rev | GATCGTAGGTGCCTTGTG |  |
| Lysozyme For | TCCCAACTCTTGACCGACGA | Is a muramidase that cleaves the linkages between N-acetylomuramid acid and N-acetylglucosamine in bacterial peptidoglycan. |
| Lysozyme Rev | AGTGGTTGCGCCATCCATAC |  |
| Cecropin D For | CTGCGCCATGTTCTTCA | Cationic antimicrobial peptide that disrupts microbial membranes, which eventually results in microbial cell death. |
| Cecropin D Rev | TCGCATCTCTGATCCTCTG |  |
| Moricin For | GCTGTACTCGCTGCACTGAT | Antibacterial activity against both Gram-positive and Gram-negative bacteria.Forms ion channels in the bacterial membrane. |
| Moricin Rev | TGGCGATCATTGCCCTCTTT |  |
| Hemolin For | CCCGAAGACGCTGGTGAATA | Functions as an opsonin that facilitates pathogen recognition and mediates hemocytic immune responses |
| Hemolin Rev | CGCACGTTCATTTGCTGTTC |  |
| GST For | GACAGAAGTCCTCCGGTCAG | Glutathione S-transferase: protect cells from oxidative stress, but they also play a central role in the detoxification of both endogenous and xenobiotic compounds. |
| GST Rev | TCCGTCTTCAAGCAAAGGCA |  |
| NOX-4 For | TGGCACGGCATCAGTTATCA | NADPH oxidase: is a pro-oxidative stress enzyme, induce oxidative stress humoral responses and secrete reactive oxygen species (ROS). |
| NOX-4 Rev | ACAGCGACTGTCATGTGGAA |  |
| NOS For | ATGAAGGTGCTGAAGTCACAA | Nitric oxide synthase: NO active the gene encoding of some antimicrobial peptides. |
| NOS Rev | GCCATTTTACAATCGCCACAA |  |
| IMPI For | ATTTGTAACGGTGGACACGA | Insect metalloproteinase inhibitor: protect AMPs against digestion by metalloproteinases secreted by the invading bacteria. |
| IMPI Rev | CGCAAATTGGTATGCATGG |  |
| Transferrin For | CCCGAAGATGAACGATCAC | Mediates nutritional immunity by sequestering iron from invading pathogens. Insects trigger a hypoferremic response after infection to limit iron availability to invading microbes. |
| Transferrin Rev | CGAAAGGCCTAGAACGTTTG |  |

**Supplementary Figure 1:** Schematic summary of the procedure used to develop and quantify the different static dual-species biofilms in microtiter plates. The *P. aeruginosa* PAO1 and PAET1 strains were detected by luciferase (*lux*) bioluminescence, and *M. abscessus* rough (R) and smooth (S) morphotypes were detected by fluorescence (GFP).

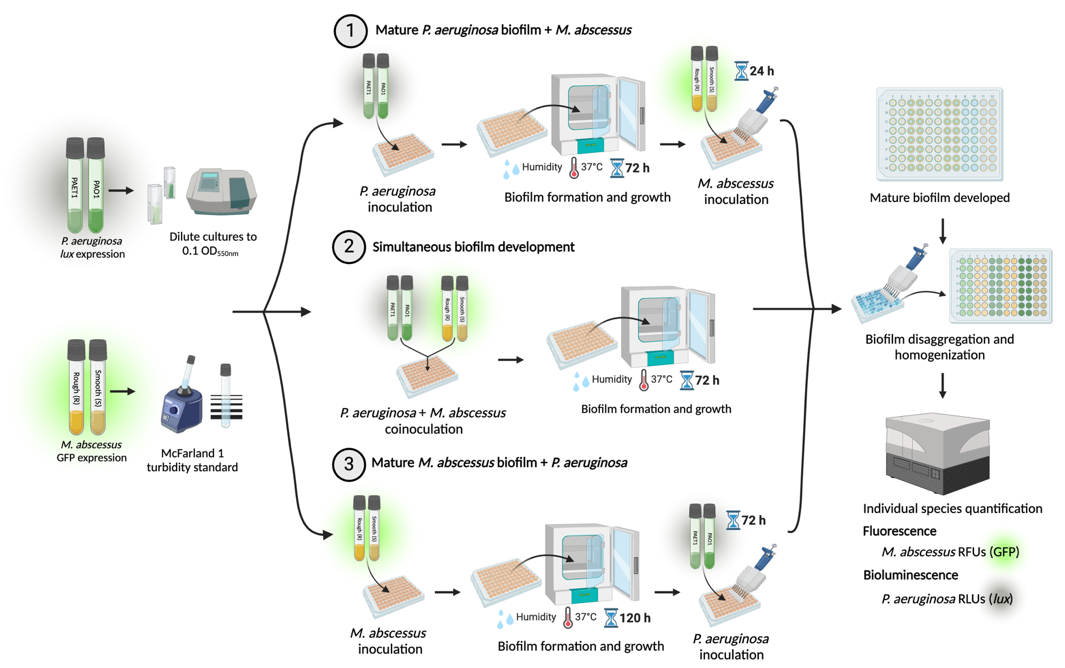

**Supplementary Figure 2:** Representative images of *P. aeruginosa* and *M. abscessus* dual-species biofilms with orthogonal projections show both bacteria’s heterogeneous spatial distribution in the biofilm. DAPI stained all cells blue, and green fluorescent protein (GFP) of plasmid pETS218 allowed mycobacterial cells to be detected in green. Scale bars correspond to 20 µm.

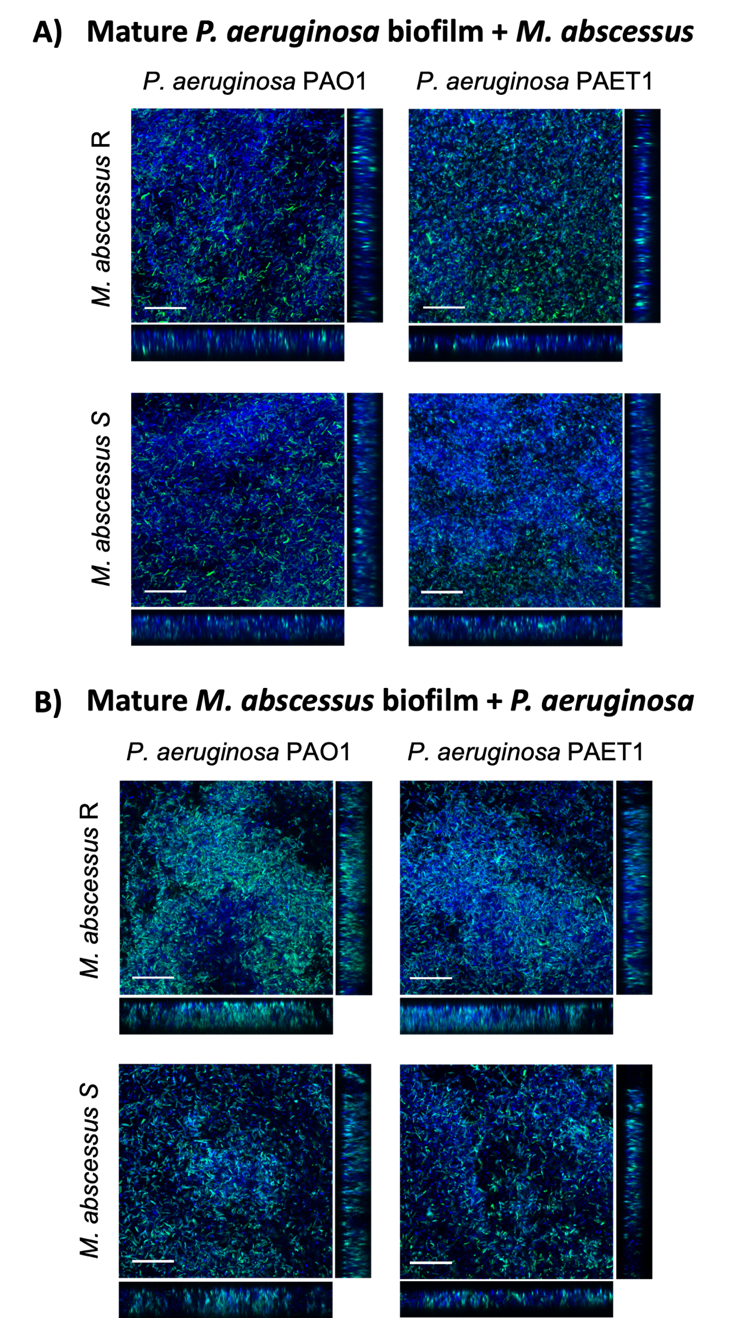

**Supplementary Figure 3: Kaplan–Meier survival curves of *Galleria mellonella* for the optimization of bacterial innocuous and lethal doses.** The strains used in this study were tested to determine doses of interest. *P. aeruginosa* was lethal at all doses tested. The doses of PAET1 strain were adjusted to obtain similar results as PAO1 (more virulent). Ten larvae were used for each bacterium and condition (n=10).

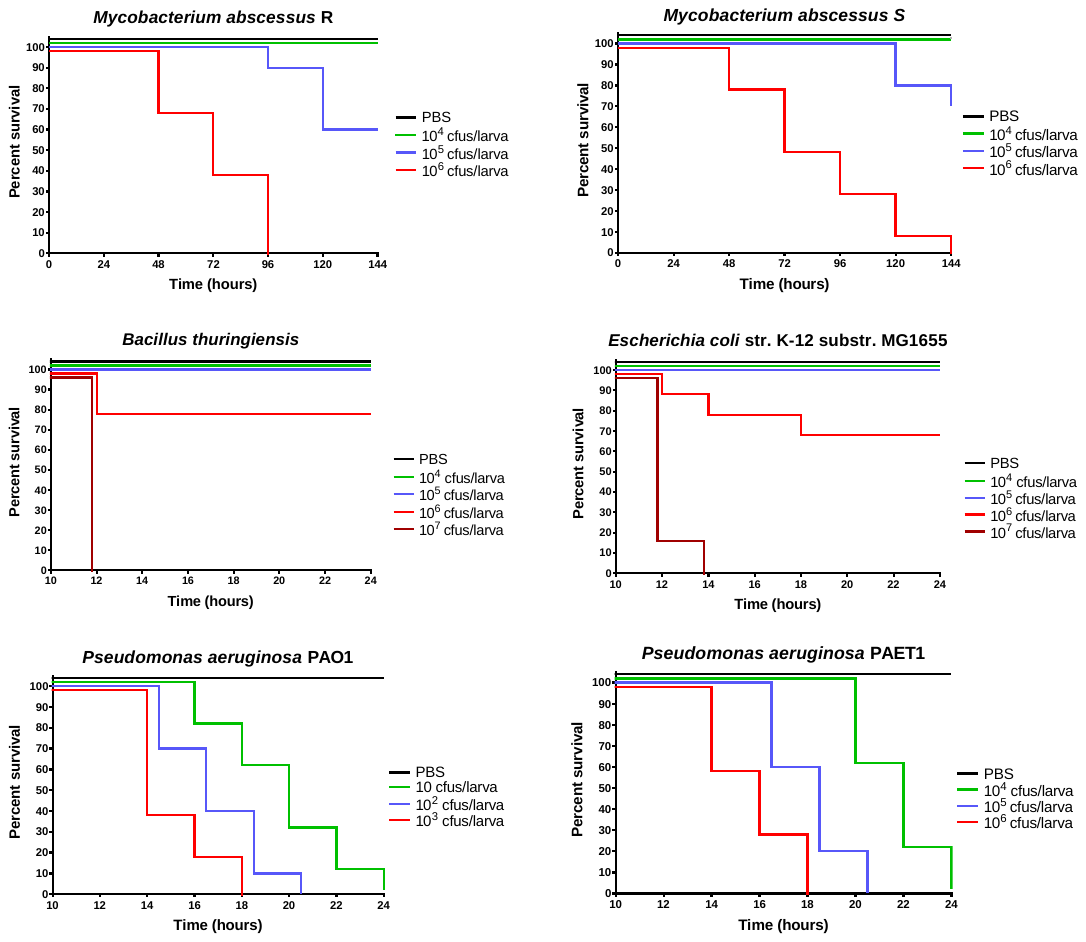

**Supplementary Figure 4: Images of *lux* expression in *G. mellonella* larvae to report *P. aeruginosa* infection.** Images were taken every 2 h until all the larvae died (-) using an Image Quant LAS 4000 (GE Healthcare, Chicago, USA) measuring chemiluminescence at a 30 sec exposure.

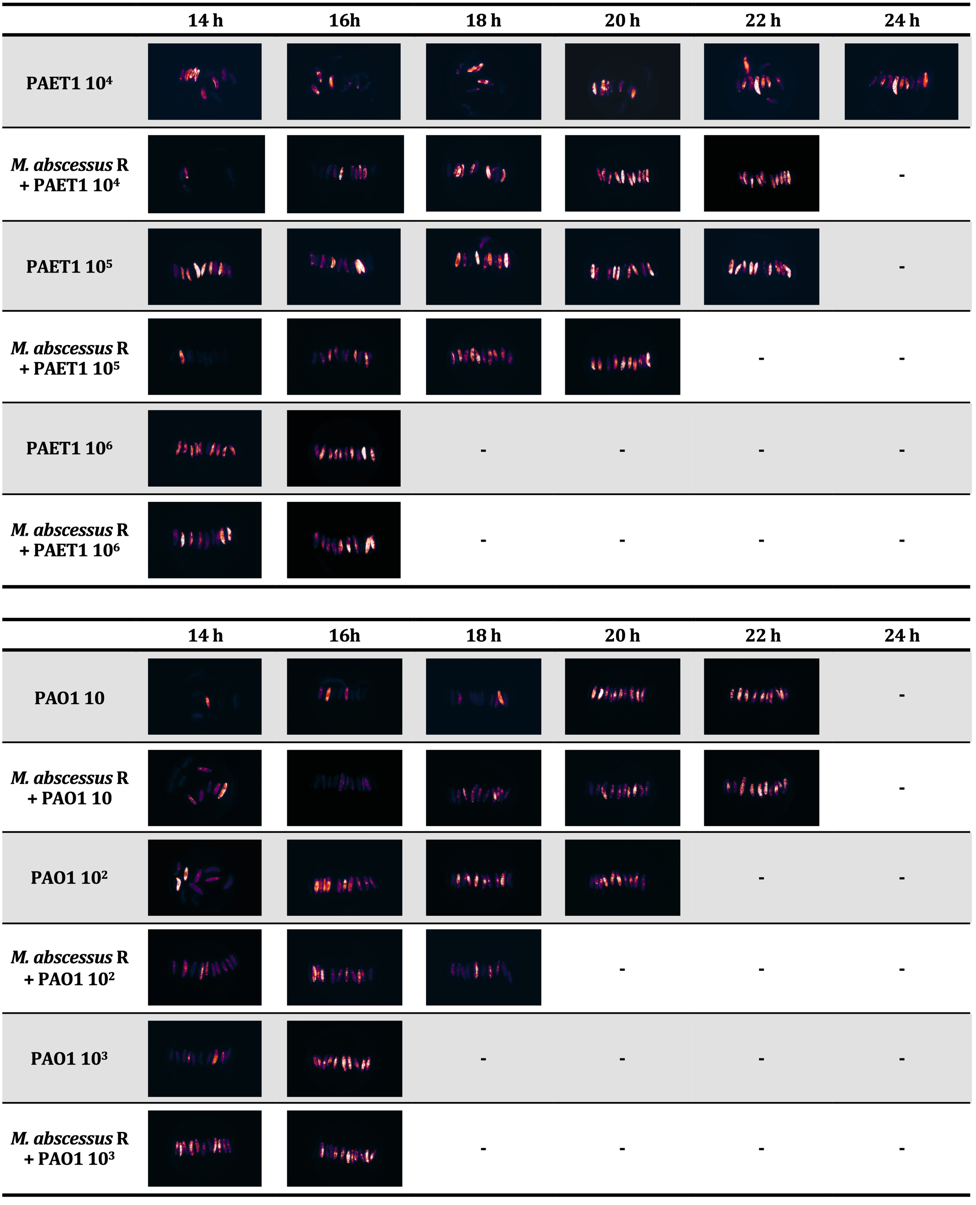

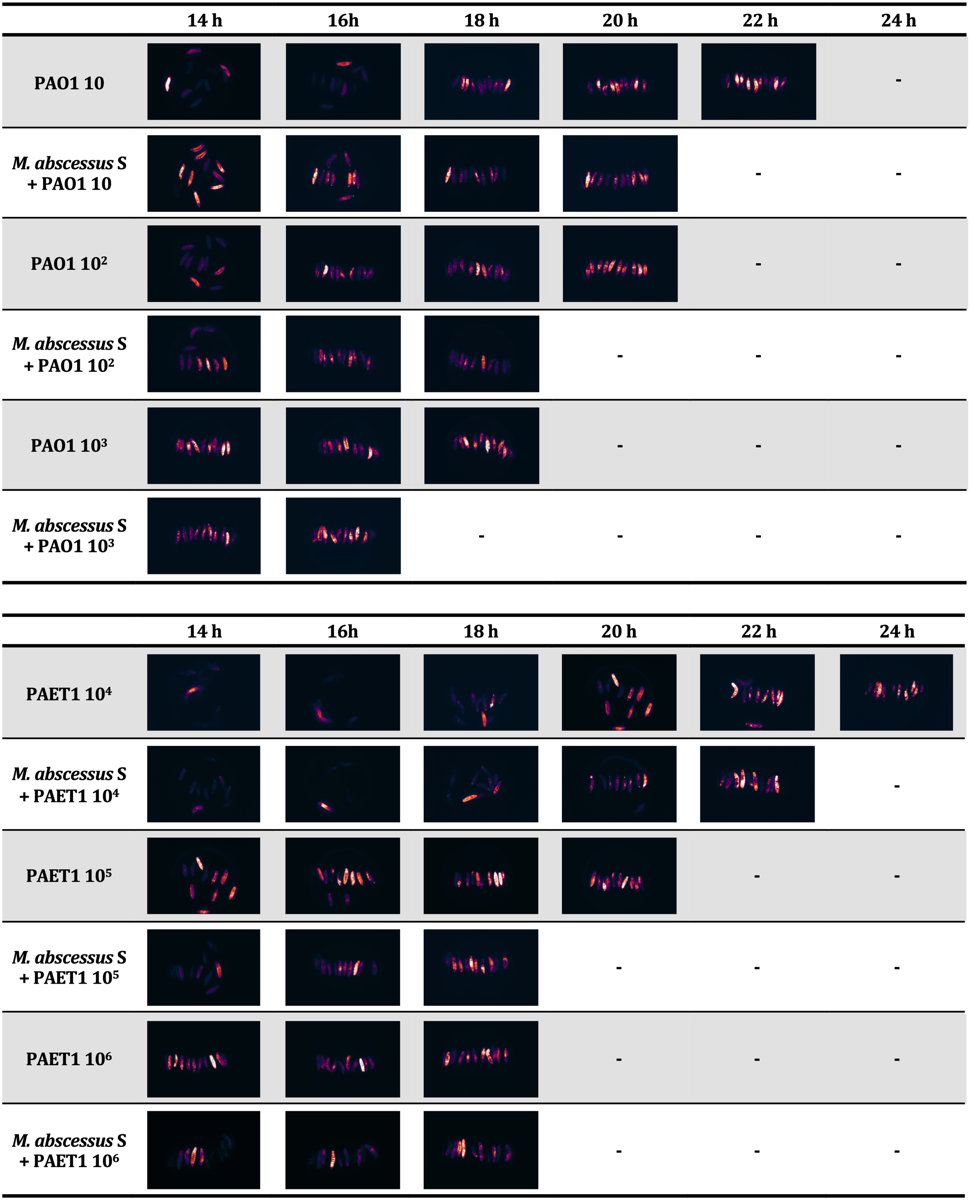

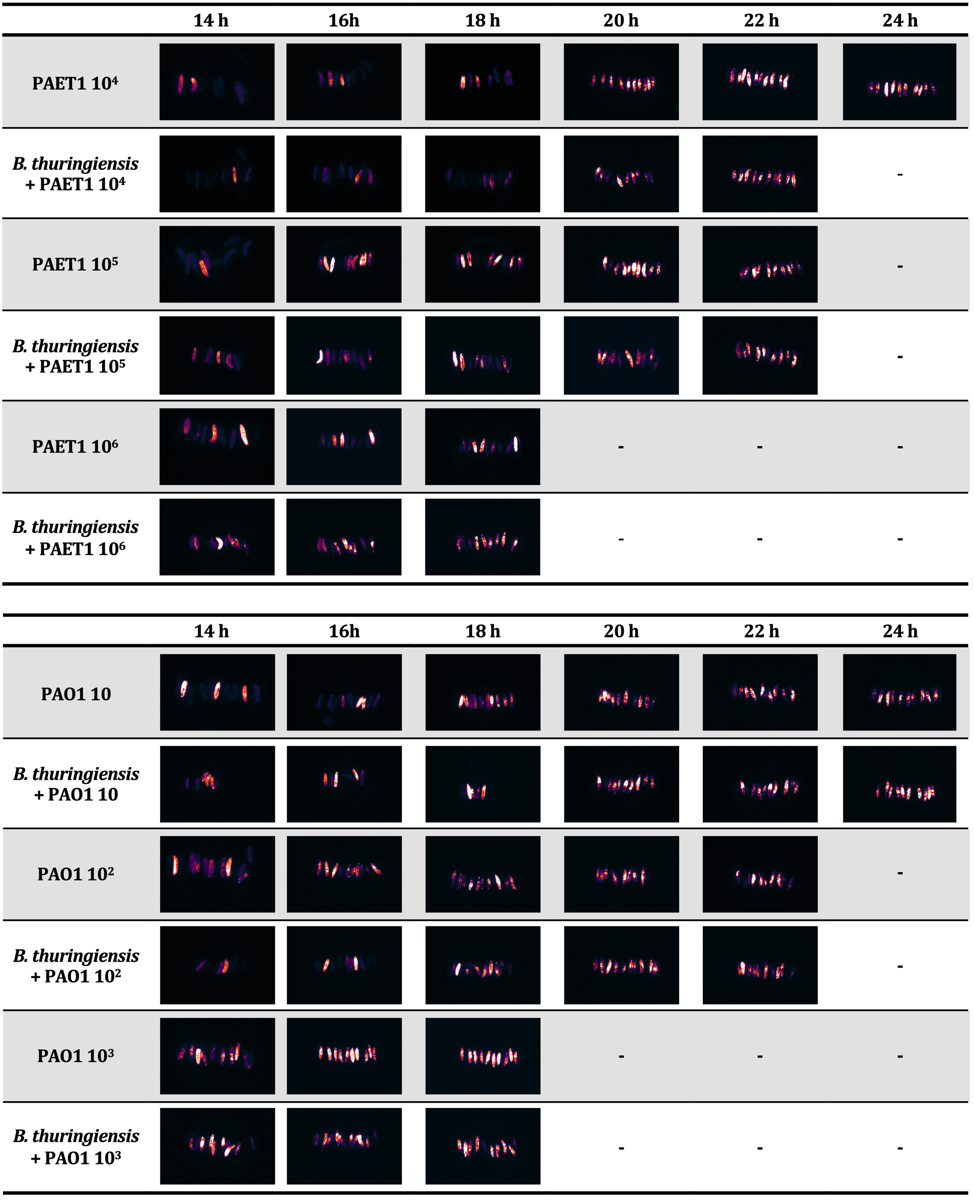

**Supplementary Figure 5:** **Direct growth inhibition assay and growth kinetics of *P. aeruginosa* strains culture with bacterial culture supernatants. A)** Images of agar plates with *M. abscessus* R and S cultures and perpendicular streaks of the different strains of *P. aeruginosa, E. coli* and *B. thuringiensis* to visualize possible inhibitory growth effects. **B)** *P. aeruginosa* PAO1 and PAET1 liquid cultures growth kinetics after adding different amounts of supernatant from cultures of *M. abscessus* R and S. **C)** *P. aeruginosa* PAO1 and PAET1 growth kinetics by adding different amounts of supernatant from cultures of *E. coli* and *B. thuringiensis*. The absorbance (OD _550 nm_) and bioluminescence (Relative Light Units, RLUs) of *P. aeruginosa* planktonic cultures are quantified in 96-well plates (100 µl/well) every 15 min in contact with different volumes of other bacterial supernatants (SN); 10, 30 and 50 µl.

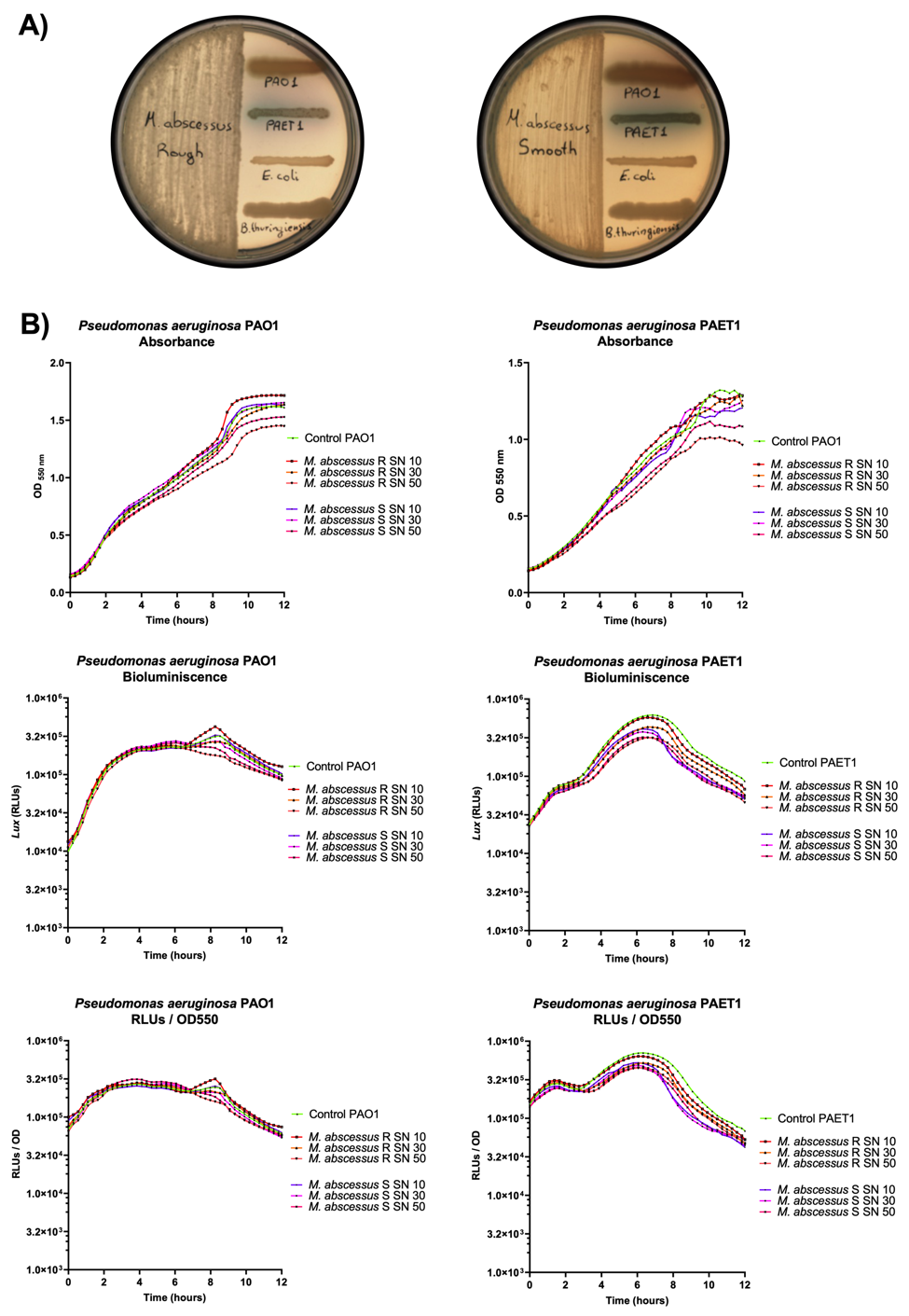

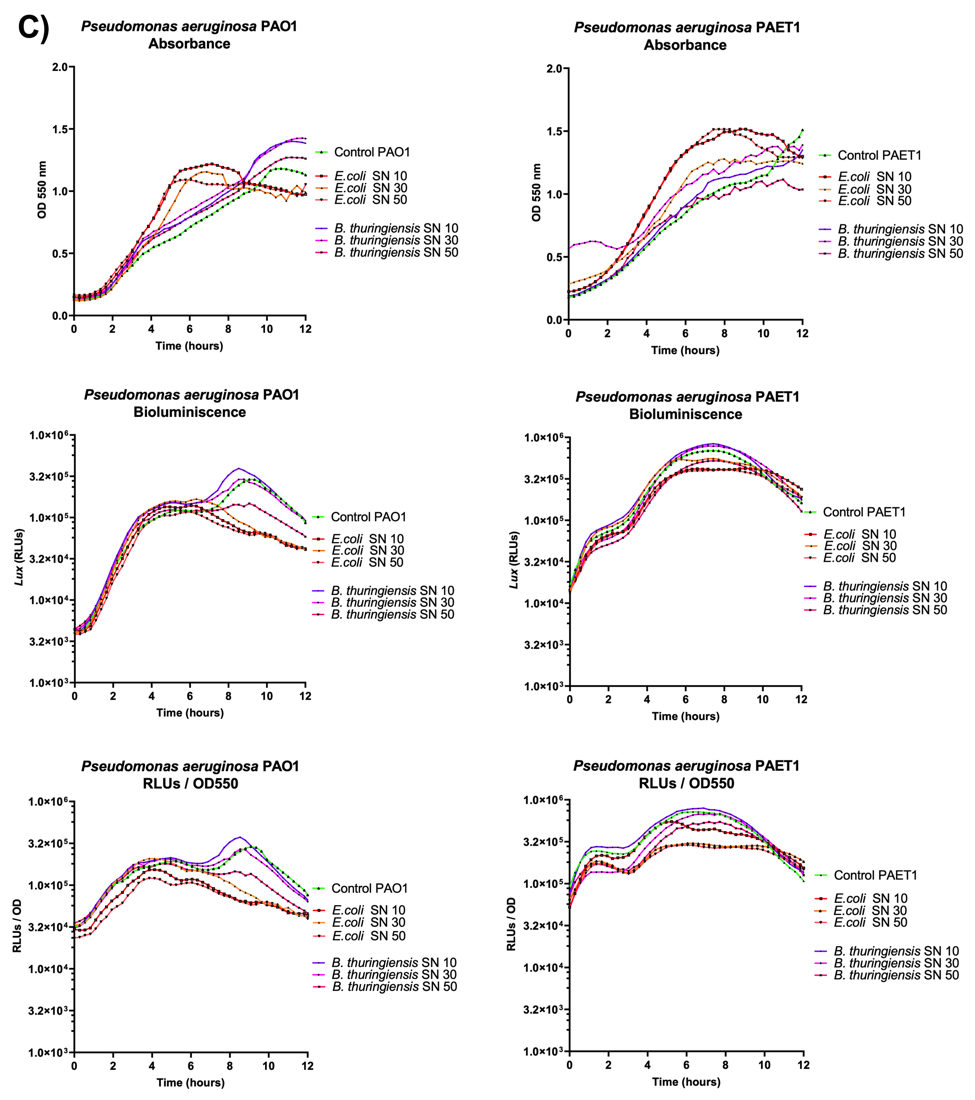

**Supplementary Figure 6: Phagocytosis of *M. abscessus* bacilli by *G. mellonella* hemocytes and prevalence inside the larvae over time.** **A)** Confocal microscope micrographs and Imaris software reconstruction of *M. abscessus* phagocytosed by *G. mellonella* hemocytes. Cell nuclei was dye with DAPI (blue), vesicular network and plasma membrane were dye with FM 4-64 (red) (N-3-Triethylammoniumpropyl-4-6-4-Diethylamino Phenyl Hexatrienyl Pyridinium Dibromide) and mycobacteria was detected through GFP (green) expression. Scale bars correspond to 10 μm in XY and 5μm in XZ reconstructions. **B)** *M. abscessus* R and S morphotypes colony forming units (cfus) recovered from *G. mellonella* hemolymph after infection over time.

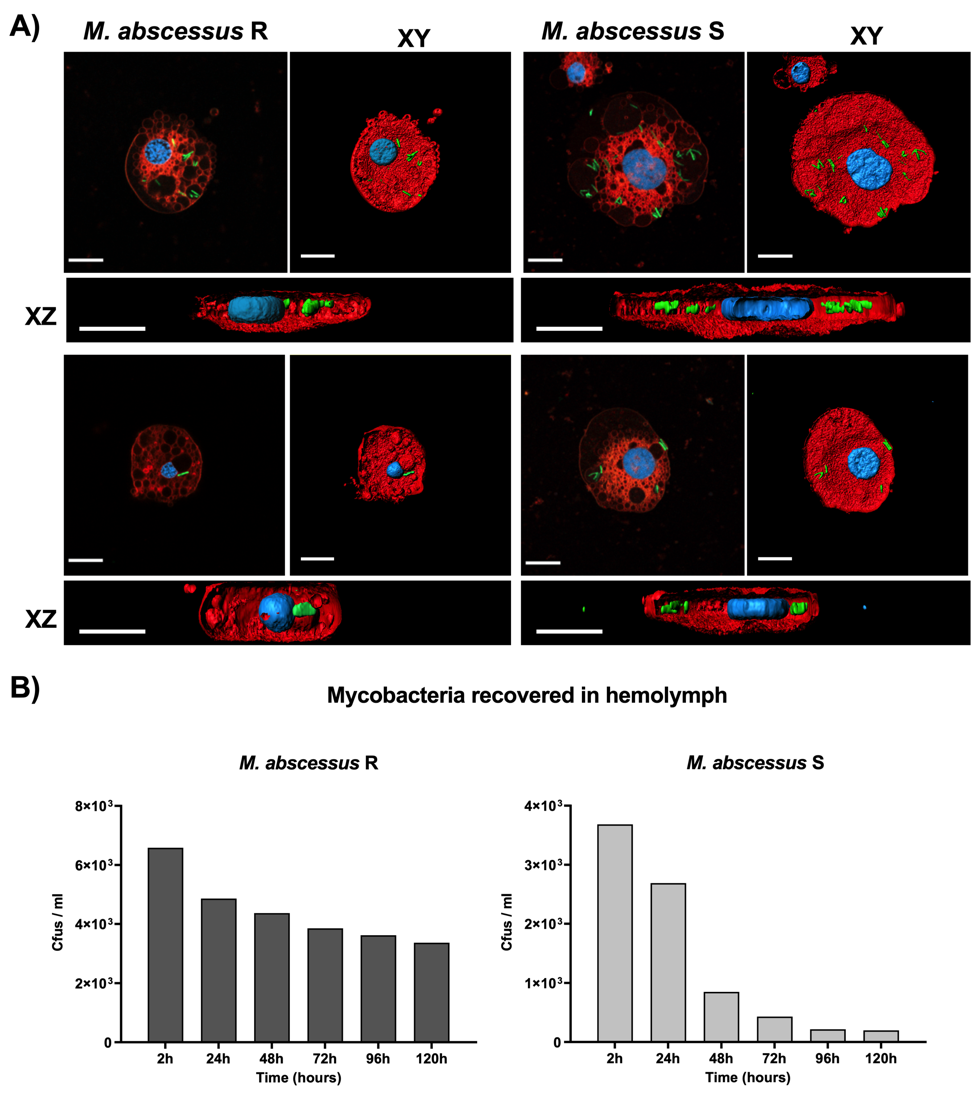
